# Mfsd2b and Spns2 are essential for maintenance of blood vessels during development and protection of anaphylaxis

**DOI:** 10.1101/2021.11.15.468739

**Authors:** Thanh Nha Uyen Le, Toan Q. Nguyen, Yen Thi Kim Nguyen, Clarissa Kai Hui Tan, Farhana Tukijan, Ludovic Couty, Zafrul Hasan, Pazhanichamy Kalailingam, Markus Wenk, Amaury Cazenave-Gassiot, Eric Camerer, Long N. Nguyen

**Affiliations:** Department of Biochemistry, Yong Loo Lin School of Medicine, National University of Singapore Singapore 119228; Université de Paris, PARCC, INSERM U970, 56 Rue Leblanc, F-75015 Paris, France; Singapore Lipidomics Incubator (SLING), Life Sciences Institute, National University of Singapore, Singapore 117456; Cardiovascular Disease Research (CVD) Programme, Yong Loo Lin School of Medicine, National University of Singapore, Singapore 117545; Immunology Program, Life Sciences Institute, National University of Singapore, Singapore 117456; Immunology Translational Research Program, Yong Loo Lin School of Medicine, National University of Singapore, Singapore 117456

**Keywords:** Sphingosine-1-phosphate, S1P transporters, Mfsd2b, Spns2

## Abstract

Sphingosine-1-phosphate (S1P) is a potent lipid mediator that is secreted by several cell types to induce signaling. We recently showed that Mfsd2b is an S1P transporter from hematopoietic cells, which contributes approximately 50% plasma S1P. To further determine the sources of plasma S1P, here, we report the characterizations of compound deletions of Mfsd2b and Spns2, another S1P transporter from endothelial cells. Global deletion of Mfsd2b and Spns2 (gDKO) resulted in embryonic lethality between E13.5 and E14.5 with severe hemorrhage that largely recapitulated the phenotypes from global S1P1 knockout mice, indicating that together with Mfsd2b, Spns2 also provides embryonic source of S1P for S1P1 stimulation. The hemorrhagic phenotypes in gDKO embryos were accompanied by increased angiogenesis and defects of tight junction proteins, indicating that S1P from Mfsd2b and Spns2 is essential for blood vessel integrity and maturation. The various sources of S1P in postnatal stages are yet to be fully understood. Postnatal ablation of S1P synthesis enzymes using Mx1Cre shows that Mx1Cre-sensitive cells provide most of plasma S1P. Interestingly, we showed that compound postnatal deletion of Mfsd2b and Spns2 using Mx1Cre (ctDKO-Mx1Cre) resulted in maximal reduction of 80% plasma S1P. Thus, a small amount of plasma S1P is supplied from other sources independent of Mfsd2b and Spns2. Nevertheless, the vasculature in the lung of ctDKO-Mx1Cre mice was compromised. Furthermore, ctDKO-Mx1Cre mice also exhibited severe susceptibility to anaphylaxis, indicating that S1P from Mfsd2b and Spns2 is indispensable during vascular stress. Together, our results show that Mfsd2b and Spns2 provide a critical source of S1P for embryonic development and they also provide a majority of plasma S1P for vascular homeostasis.

## Introduction

Sphingosine-1-phosphate (S1P) is a potent lipid mediator that is required for lymphocyte egress from lymphoid organs and essential for blood vessel development and function. S1P exerts signaling roles via 5 cognate S1P receptors (S1P1-5). In cells, S1P is synthesized from sphingosines by sphingosine kinase 1 and 2 (SphK1 and 2) (Yanagida and Hla, 2017). Extracellular SphK1 can also generate S1P, but it remains to be determined whether this pathway is physiologically relevant (Venkataraman et al., 2006). S1P is a polar lipid. Therefore, its transport across the lipid bilayers must be mediated by transporters. The concentrations of extracellular S1P are tightly regulated by multiple enzymes and transporters to form a gradient with levels ranging from <1nM in interstitial fluid to ~100nM in lymph and ~1000nM in blood (Hla and Dannenberg, 2012), (Spiegel and Milstien, 2011), (Rosen and Goetzl, 2005). Lymphocytes rely on the gradient of S1P for trafficking. Whether vascular cells are also sensitive to S1P gradients remains to be established.

An important role of S1P signaling is the regulation of lymphocyte trafficking. T and B lymphocytes from lymphoid tissues sense S1P that is released from lymphatic endothelial cells for egress (Yanagida and Hla, 2017), (Schwab and Cyster, 2007), (Baeyens and Schwab, 2020). Mechanistically, S1P binds and stimulates S1P1 on the plasma membrane of lymphocytes that drives their migration from lymph node parenchyma to lymphatic circulation. Desensitization of S1P1 with Fingolimod and other S1P modulators to block lymphocyte egress has been employed for treatment of multiple sclerosis (Rosen et al., 2009), (Chun et al., 2019), (Chun et al., 2021). S1P signaling also plays essential roles in the vasculature. Several S1P receptors including S1P1, S1P2, and S1P3 are expressed in different cell types in blood vessels (Yanagida et al., 2020). Deletion of S1P1 results in embryonic death from E12.5 and E14.5 with severe hemorrhage (Liu et al., 2000). Specific ablation of endothelial S1P1 in adult mice also results in impaired vascular integrity (Jung et al., 2012), (Gaengel and Betsholtz, 2013), (Yanagida et al., 2017). Compound deletion of S1P1 with either S1P2 or S1P3 or triple knockout of these receptors results in earlier lethality reminiscent of compound deletion of sphingosine kinase 1&2 (Kono et al., 2004), (Mizugishi et al., 2005) emphasizing that they play non-redundant roles in blood vessels. Mechanistically, S1P signaling is employed to stabilize blood vessels by reducing VEGF-driven angiogenesis and increasing the expression of tight junction proteins. It also regulates blood vessel contraction via S1P1-dependent nitric oxide release in endothelial cells and constriction via S1P2&3 in smooth muscle cells. In line with essential roles of S1P signaling in blood vessels, compound conditional deletion of SphK1 and 2 results in vascular leakage and sensitization of vascular insults (Camerer et al., 2009), (Gazit et al., 2016).

In order to understand how S1P signaling is regulated it is important to determine which cell types release S1P. By selective deletion of SphK1 and SphK2, it has been established that endothelial and hematopoietic cells are the major sources of plasma S1P (Gazit et al., 2016). In line with these studies, it has been shown that Spns2 exports S1P from the endothelial cells, whereas Mfsd2b exports from erythrocytes and platelets, respectively (Fukuhara et al., 2012), (Vu et al., 2017), (Chandrakanthan et al., 2021). While Spns2 deletion results in strong lymphopenia due to a profound decrease in lymph S1P, vascular development and function are normal in Spns2 knockout mice (Mendoza et al., 2012). Global deletion of Mfsd2b results in approximately 50% reduction of plasma S1P and renders mice susceptible to passive systemic anaphylaxis (Vu et al., 2017). However, these isolated knockout mice thrive to adulthood, suggesting that Mfsd2b and Spns2 provide redundant sources of S1P to maintain vascular development and homeostasis. Several ABC transporters have also been reported to transport S1P (Mitra et al., 2006), (Kobayashi et al., 2009). To understand the roles of Mfsd2b and Spns2 in S1P export and to address if additional S1P transporters exist, we generated compound deletions of Mfsd2b and Spns2 before and after birth. Our results show that Mfsd2b and Spns2 are essential for embryonic development. Lack of both genes resulted in lethality with hemorrhagic phenotypes analogous to the S1P1 knockout mice, suggesting that they probably deliver S1P to endothelial S1P1 for signaling. Furthermore, we show that Mfsd2b and Spns2 coordinate to provide a major proportion of plasma S1P to maintain the vasculature in adulthood. Intriguingly, our results suggest that there may be a novel source of S1P that is independent from Mfsd2b and Spns2 from Mx1Cre sensitive cells.

## Results

### Global deletion of Mfsd2b and Spns2 causes embryonic lethality

We and others have recently shown that Mfsd2b and Spns2 are two S1P exporters in hematopoietic and vascular systems, respectively (Fukuhara et al., 2012), (Vu et al., 2017), (Mendoza et al., 2012). Global deletion of either Mfsd2b or Spns2 results in incomplete reduction of plasma S1P and there is no overt embryonic phenotypes (Fukuhara et al., 2012), (Vu et al., 2017). Single cell RNAseq showed that Mfsd2b is specifically expressed in primitive and definitive red blood cells (RBC) (Cao et al., 2019) (**Supplemental Figure 1A, B**). Interestingly, Spns2 expression is also found in primitive RBC and endothelial cells (Cao et al., 2019) (**Supplemental Figure 1C**). To examine whether Mfsd2b and Spns2 are functional during embryonic development, we isolated primitive RBC and fetal liver cells the corresponding knockout embryos for transport assays. We showed that primitive RBC from Mfsd2b^-/-^ embryos exhibited approximately 50% reduction of S1P transport activity (**Figure 1A**). Fetal liver cells from Mfsd2b^-/-^ embryos also exhibited slightly reduced S1P transport activity (**Figure 1B**). Surprisingly, primitive RBC, but not fetal liver cells from Spns2^-/-^ embryos also had slightly, but significant reduction of S1P transport activity (**Figure 1C-D**). These data suggest that Mfsd2b and Spns2 transport functions are active in primitive RBC during fetal development. To examine if they are the only physiological S1P transporters, we generated the global knockout of both Mfsd2b and Spns2 in mice (Mfsd2b^-/-^;Spns2^-/-^, hereafter gDKO) by intercrossing Mfsd2b^-/-^;Spns2^+/-^ mice. Although these male and female breeders carrying a single allele of Spns2 were healthy and fertile, we observed a small litter size, suggestive of embryonic lethality. We found that gDKO embryos were viable at embryonic day 12.5 (E12.5) with little obvious defects, although blood vessels in the lower dorsal parts were dilated (**Figure 1E, arrowheads**). However, viable gDKO embryos exhibiting noticeable signs of hemorrhage were found at E13.5 (**Figure 1E, arrows; Figure 1F, arrowhead;** and **Supplemental Figure 2**). We found dead gDKO embryos with signs of severe hemorrhage, especially in the lower dorsal parts, at E14.5 (**Figure 1E, arrows**). These results indicate that gDKO embryos die between E13.5 and E14.5 (**Table S1**). The observed hemorrhagic phenotypes of gDKO embryos are largely reminiscent of S1P1 knockout embryos, but slightly less severe compared to double knockout of sphingosine kinases (SphK1/2) (Liu et al., 2000), (Mizugishi et al., 2005). Thus, our data show that the combined source of S1P from Mfsd2b and Spns2 is critical for survival and that they are likely the sole S1P transporters during embryonic development.

**Figure 1.**
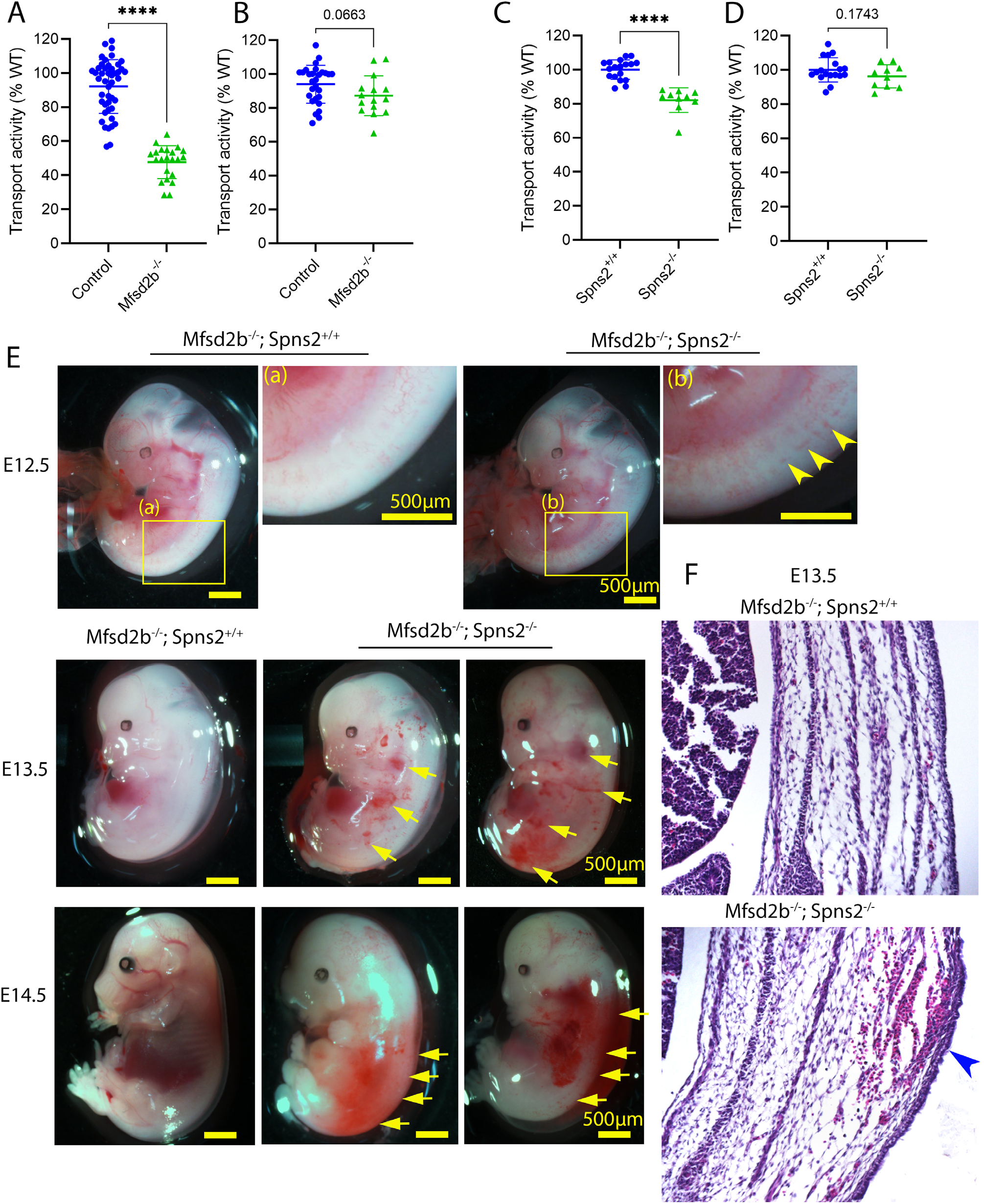
Compound deletion of Mfsd2b and Spns2 causes early embryonic lethality. **A-B**, Transport activity of primitive RBCs (A) and fetal liver cells (B) from Mfsd2b^-/-^ and control embryos. **C-D**, Transport activity of primitive RBCs (C) and fetal liver cells (D) from Spns2^-/-^ and wild-type embryos, respectively. Primitive RBCs and fetal liver cells from E17.5-18.5 Mfsd2b^-/-^ and Spns2^-/-^ embryos were collected for transport activity. Transport activity of primitive RBC from Mfsd2b^-/-^ or Spns2^-/-^ embryos were normalized to that of control littermates (WT and heterozygous embryos). Each symbol represents an embryo. ****P<0.0001; t-test. **A**, Deletion of Mfsd2b and Spns2 results in lethality between E13.5 and E14.5. In comparison with Mfsd2b knockout embryos carrying native Spns2 alleles used as controls, embryos with complete loss of both Mfsd2b and Spns2 genes (gDKO) were viable and had normal morphology at E12.5. Blood vessels from lower body of gDKO embryos were slightly dilated at E12.5 (top panel, arrowheads) and became prominent at E13.5 (middle panel, arrows). Non-viable gDKO embryos with severe hemorrhage were observed at E14.5 (low panel, arrows). gDKO embryos died between E13.5 and E14.5. The experiments were performed multiple times as shown in Table S1; (a) and (b), enlarged images of lower dorsal regions of control and gDKO embryos. **B**, Representative H&E images of a transverse section of E13.5 control and gDKO embryos. Arrowhead shows the bleeding in the dermis. Images were taken with 20X objective.

### Embryos lacking Mfsd2b and Spns2 exhibit hemorrhages and vascular defects

To gain insights into the hemorrhagic phenotype, especially in the developing brain, we stained brain sections of E13.5 embryos with vascular markers such as CD31, Glut1, and red blood cells (RBCs) marker Ter119. The vascular density and growth in the brain of gDKO appeared normal in all regions (**Figure 2 and Supplemental Figure 3A**). However, in addition to peripheral hemorrhage as shown in **Figure 1**, we found that blood vessels in the central nervous system (CNS) of E13.5 gDKO embryos were ruptured with extravasation of RBCs in the ganglionic eminences and hindbrain/hypothalamus (**Figure 2A, B, arrowheads** and **Supplemental Figure 3B, C**). RBCs were detected in the brain parenchyma, indicating impaired integrity of CNS blood vessels of gDKO (**Figure 2C** and **Supplemental Figure 3B, C**). Interestingly, deletion of S1P1 results in lethality between E12.5 and E14.5 with cerebral bleeding, accompanied by increased angiogenesis (Liu et al., 2000), (Mizugishi et al., 2005). We also found that CNS blood vessels were slightly dilated in the ganglionic eminences, but not in the midbrain of gDKO embryos (**Figure 2D, E**). Blood vessels near the ventricles were severely dilated, a phenotype that is similar to micro-aneurisms (**Figure 2A, B, arrows**). Our results show that deletion of Mfsd2b and Spns2 causes both peripheral and cerebral hemorrhage during embryonic development.

**Figure 2.**
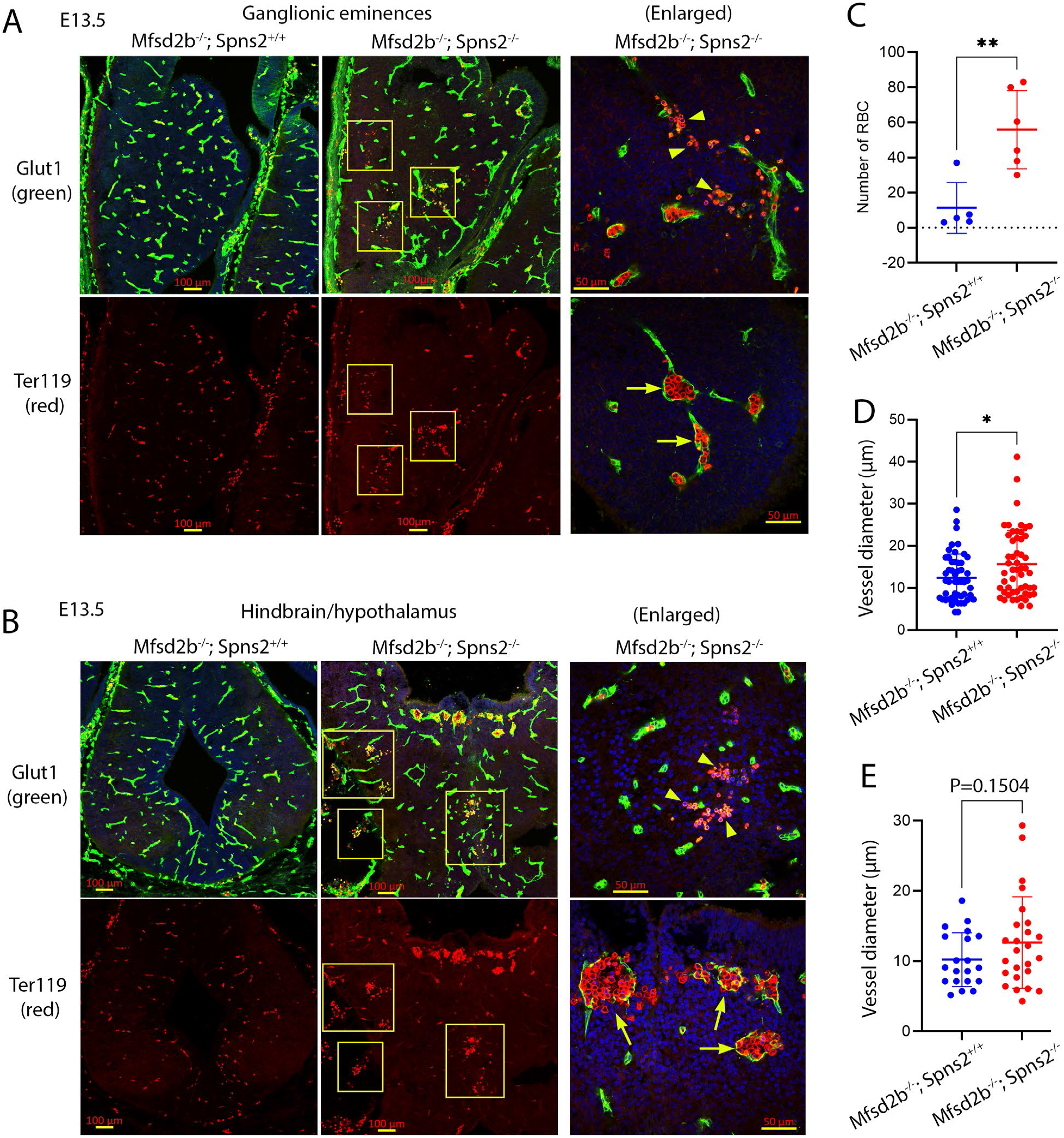
Global deletion of Mfsd2b and Spns2 results in cerebral hemorrhage and vascular defects. **A,** Representative images of ganglionic eminence (GE) regions of E13.5 Mfsd2b^-/-^ (control littermates) and gDKO embryos. gDKO embryos displayed hemorrhage (boxes and arrowheads) and dilated blood vessels (arrows); n=6 per genotype. **B,** Hemorrhage in hindbrain regions of E13.5 gDKO embryos (boxes show hemorrhagic regions). The global double knockout embryos also exhibited local hemorrhage near ventricular regions (arrowheads). Cerebral blood vessels of E13.5 gDKO fetuses were also dilated (arrows). **C**, Number of RBCs present in brain parenchyma of gDKO embryos in GE regions. Each symbol represents one field from one embryo (n=5 for controls and n=6 for gDKO). Results are mean and SD, **P<0.01; t-test. **D-E,** Diameter of blood vessels in GE (D) and midbrain (E) regions of E13.5 Mfsd2b^-/-^ and gDKO. Significant increase in diameter of blood vessels in GE regions of gDKO embryos. Each symbol represents one vessel from 5-6 embryos per genotype. Results are mean and SD, *P<0.05; t-test.

### Lack of embryonic sources of S1P from Mfsd2b and Spns2 results in cerebral blood vessel rupture

The hemorrhagic phenotypes observed in the body and brain of gDKO embryos suggest that lack of S1P secretion from Mfsd2b and Spns2 is detrimental to blood vessel integrity in the brain. First, we tested whether the coverage of pericytes, the mural cells of CNS blood vessels, is reduced in E13.5 gDKO blood vessels. Our results showed that pericyte coverage shown by PDGFRβ expression in CNS blood vessels of gDKO embryos was comparable with littermate controls (**Figure 3A**). The expression of N-cadherin was also unaffected in the brain of gDKO embryos (**Supplemental Figure 4A**). Disruption of TGFβ or canonical WNT signaling pathways results in brain hemorrhage (Stenman et al., 2008), (Daneman et al., 2009), (Wang et al., 2018), (Li et al., 2011), (Arnold et al., 2014), (Dave et al., 2018), (Crist et al., 2019). Therefore, we next examined if expression of phosphorylated Smad4 and β-catenin, the respective components of TGFβ and canonical WNT signaling pathways is altered. However, both pathways were intact in the brain of gDKO embryos (**Figure 3B**). Upregulation of PLVAP which is indicative of increased transcytosis in the blood-brain barrier of WNT signaling knockout mice was comparable between the genotypes (**Figure 3C and Supplemental Figure 4B**) (Cho et al., 2017), (Zhou et al., 2014). These data rule out the involvement of TGFβ and canonical WNT signaling in the hemorrhagic phenotypes. Lack of S1P signaling results in compromised expression of adherens and tight junction proteins that may be linked to the fragility of blood vessels in gDKO embryos. We found that the expression pattern of Claudin-5 was disrupted in the hemorrhagic vessels (**Figure 3D, arrowheads** and **Supplemental Figure 4C**). Additionally, expression of VE-cadherin, an adherens junction protein which is required for maintenance of junctions between endothelial cells, was also reduced in the brain of gDKO embryos (**Figure 3E,** quantification in **Supplemental Figure 4D**). S1P signaling is mediated by the activation of MAP kinases (Jo et al., 2005). We found that lack of Mfsd2b and Spns2 reduced the expression of phosphorylated ERK1/2 (p-ERK1/2), suggestive of low levels of S1P in the brain parenchyma of gDKO embryos (**Figure 3F,** quantification in **Supplemental Figure 4E**). Together, our results show that S1P released via Mfsd2b and Spns2 is required for protection of CNS blood vessel integrity during embryonic development.

**Figure 3.**
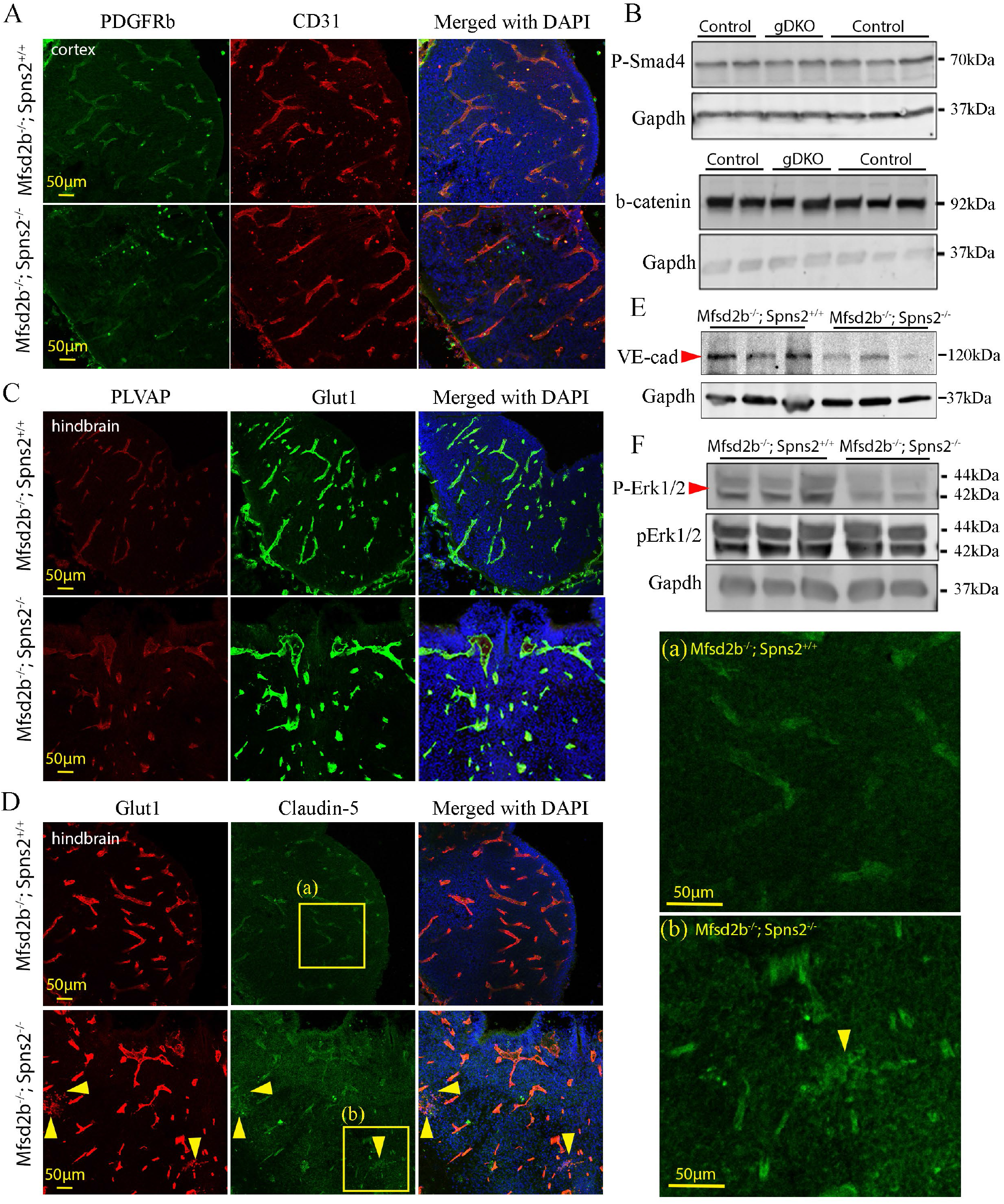
Lack of Mfsd2b and Spns2 results in local rupture of CNS blood vessels during development. **A,** Pericyte coverage was not reduced in CNS blood vessels of E13.5 gDKO embryos; n=3 per genotype. **B,** Expression of canonical WNT (β-catenin) and TGFβ signaling (phosphorylated Smad4) proteins was comparable between E13.5 controls (Mfsd2b^-/-^;Spns2^+/-^ or Mfsd2b^-/-^;Spns2^+/+^) and E13.5 gDKO. n=5 for controls and n=2 for gDKO. **C**, Immunostaining of PLVAP in CNS blood vessels of E13.5 gDKO embryos. PLVAP expression was not induced in the blood vessels of gDKO embryos compared to control; n=3 per genotype. **D**, Localization of Claudin-5 was disrupted in the ruptured cerebral blood vessels of E13.5 gDKO embryos. Shown in (a) and (b) are enlarged regions with Claudin-5 staining in the blood vessels of control and gDKO embryos, respectively. Arrowheads indicate the disrupted localization of Claudin-5; n=3 per genotype. **E**, Reduced expression of VE-cadherin (CDH5) from the whole brain lysates of E13.5 gDKO compared to controls. n=3 per genotype. **F**, Reduced expression of phosphorylated ERK1/2 from the whole brain lysates of E13.5 gDKO compared to controls. n=3 for control and n=2 for gDKO embryos.

### Embryonic sources of S1P from Mfsd2b and Spns2 are also essential for normal development of aorta

S1P signaling is required for maturation and stabilization of blood vessels (Jung et al., 2012), (Ben Shoham et al., 2012), (Gaengel et al., 2012), (Paik et al., 2004). Therefore, we examined if the development of descending aorta is affected in gDKO embryos. We found that the aortae of E13.5 gDKO embryos were deformed, in which several aortae were found with flattened structures (**Figure 4A**). Nevertheless, they had normal aortic circumferences compared to controls (**Figure 4B, C**). Interestingly, we found that the endothelial cell layer of the aorta shown by staining with CD31 exhibited increased proliferation (**Figure 4A**, **arrowheads**; **Supplemental Figure 5A, B**). The coverage of smooth muscle cells is critical for maintenance of aortic structure. We found that expression of smooth muscle cell marker αSMA surrounding aorta in the dorsal side was normal, but it was significantly thinner in the ventral side (**Figure 4D, E**). Furthermore, we found that the smooth muscle cell layer was less compacted with increased sizes of nuclei (**Figure 4F, left panel, arrowheads**). The arrangement of endothelial cells was also disorganized (**Figure 4F, arrows**) in which the expression pattern of Claudin-5 in the endothelial cells of the aorta was disrupted (**Figure 4F, right panel, arrowheads**). These results are consistent with the increased angiogenic sprouting reported in the developing aorta of global S1P1 knockout embryos (Kono et al., 2004), (Gaengel et al., 2012), although slightly less severe for the same developmental stage and demonstrate that Mfsd2b and Spns2 are required for S1P delivery to maintain normal aortic structures.

**Figure 4.**
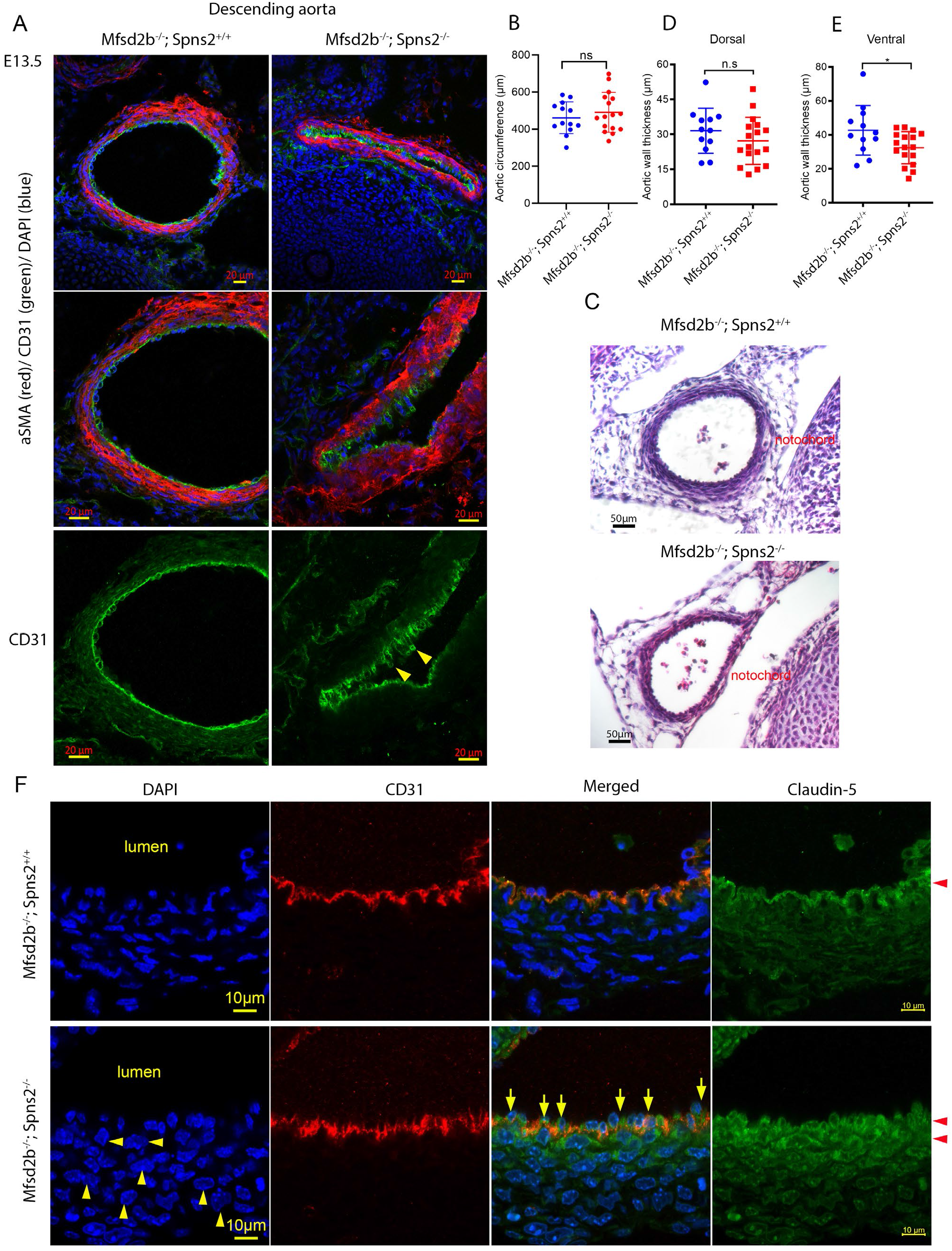
Defects in aortic structures in gDKO embryos. **A,** Aortae of gDKO embryos were deformed. Representative images of the localization of smooth muscle cells (αSMA, red) and endothelial cells (CD31, green) in the descending aorta of E13.5 control (Mfsd2b^-/-^) and gDKO embryos. **B**, Measurement of circumference of aortae from control (Mfsd2b^-/-^) and gDKO embryos. Each symbol represents one aorta from 9-10 embryos per genotype. Results are mean and SD; ns, not significant. **C**, Representative H&E images of aorta from control and gDKO embryos. **D-E**, Thickness of αSMA layer from dorsal (D) and ventral (E) sides. Ventral αSMA layer was reduced in aortae of E13.5 gDKO embryos compared to controls. Each symbol represents one aorta from 9-10 embryos per genotype. Results are mean and SD, *P<0.05; t-test. **F**, Defects in endothelial layer of aortae of E13.5 gDKO embryos (arrows). Expression of Claudin-5 was also found ectopically in the vascular smooth muscle cell layer (arrowheads). Nuclei of smooth muscle cells in aortae of E13.5 gDKO embryos exhibited round structures (left panel, arrowheads) compared to condensed structures of controls. Experiments were performed from 4-5 embryos per genotype.

### Mfsd2b and Spns2 from Mx1Cre-sensitive cells constitute a major source of plasma S1P

Previous studies have shown that conditional deletion of SphK1&2 using Mx1Cre resulted in >90% S1P reduction (Gazit et al., 2016). As a consequence, these knockout mice exhibited severe pulmonary leakage at basal condition (Gazit et al., 2016). Mx1Cre drives the deletion of floxed genes in hematopoietic cells, endothelial cells, and hepatocytes suggesting that these cells are the major sources of plasma S1P (Kuhn et al., 1995). To determine, if Mfsd2b and Spns2 mediate the export of S1P derived from SphK1&2 in these cells, we generated compound conditional deletion of Mfsd2b and Spns2 using Mx1Cre (Mfsd2b^f/f^;Spns2^f/f^;Mx1Cre, hereafter ctDKO-Mx1Cre). First, to address that Mx1Cre mediated efficient deletion of Mfsd2b and Spns2 in ctDKO-Mx1Cre mice, we measured S1P transport activity in erythrocytes (RBCs) and quantified numbers of white blood cells (WBCs), two phenotypes that are attributed to the loss of Mfsd2b and Spns2, respectively. Indeed, S1P transport activity in RBCs isolated from ctDKO-Mx1Cre showed similar defects to those seen in RBCs from global Mfsd2b, but not in RBCs from Spns2 knockout mice (**Figure 5A, B**). Expression of Mfsd2b protein in RBCs isolated from ctDKO-Mx1Cre mice was also abolished (**Figure 5C**). Deletion of Mfsd2b increases erythrocyte cell volume. Indeed, this is recapitulated in erythrocytes from ctDKO-Mx1Cre mice (**Figure 5D**). In line with the efficient deletion of Spns2, peripheral WBC count from ctDKO-Mx1Cre mice was as low as in global Spns2 KO mice (**Figure 5E**). These data indicate that expression of Mfsd2b and Spns2 was abolished in ctDKO-Mx1Cre mice. Second, we analyzed plasma S1P collected from ctDKO-Mx1Cre mice at 1 and 4 months post-injection of polyIC (**Figure 5F**). In these analyses, global deletion of Mfsd2b (Mfsd2b^-/-^ mice) resulted in ~55% reduction of plasma S1P, whereas deletion of Spns2 (Spns2^-/-^ mice) caused ~12% reduction of plasma S1P (**Figure 5G-I**). Plasma S1P levels in ctDKO-Mx1Cre was decreased by ~70% at 1 month after polyIC treatment (**Figure 5G**). Notably, plasma S1P levels in ctDKO-Mx1Cre mice were reduced by ~80% at 4 months after polyIC injection (**Figure 5I**). Collectively, these results indicate that both Mfsd2b and Spns2 coordinate to contribute to plasma S1P and that these two transporters from Mx1Cre-sensitive cells provide approximately 80% plasma S1P. Our results also suggest that additional source of >10% S1P are present among Mx1Cre positive cells. Based on these results, we selected ctDKO-Mx1Cre mice after >3 months injection with polyIC and their WBC count is lower than 2500 cells/μl for subsequent experiments.

**Figure 5.**
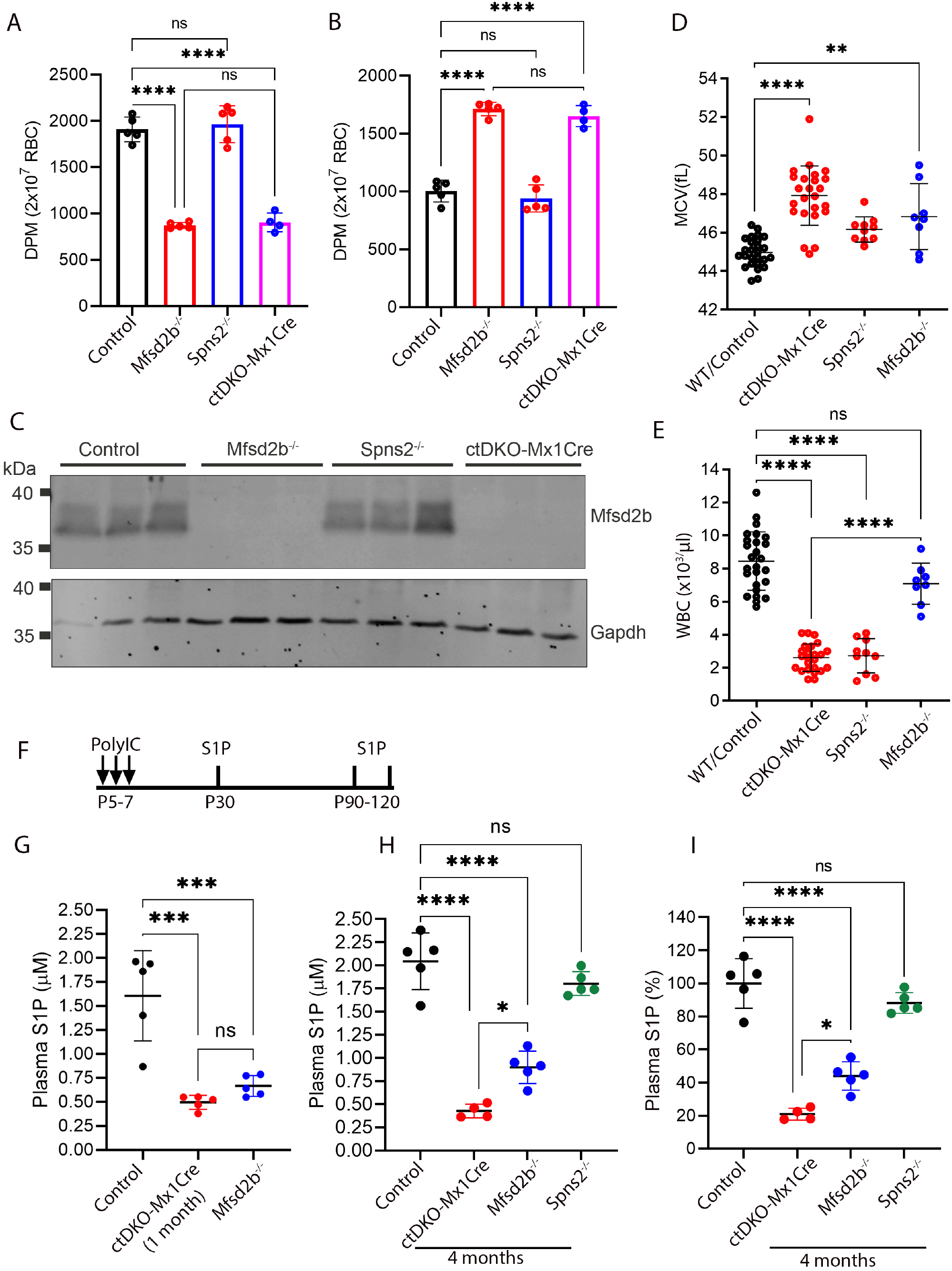
Mfsd2b and Spns2 coordinate to supply most of plasma S1P. **A-B,** Reduced S1P transport activity of erythrocytes from ctDKO-Mx1Cre mice. Extracellular (A) and intracellular (B) S1P levels from erythrocytes from control, Mfsd2b^-/-^, Spns2^-/-^, and ctDKO-Mx1Cre mice. Transport activity of erythrocytes from ctDKO-Mx1Cre mice were identical to that of erythrocytes from Mfsd2b^-/-^ mice. Data are mean and SD. Each symbol represents one mouse (n=4-5 per genotype). **C,** Western blot analysis of Mfsd2b expression in erythrocytes from control, Mfsd2b^-/-^, Spns2^-/-^, and ctDKO-Mx1Cre mice. These data showed efficient deletion of Mfsd2b in erythrocytes from ctDKO-Mx1Cre mice. Experiments were performed twice with n=3. **D**, Increased MCV of erythrocytes from ctDKO-Mx1Cre mice. **E,** White blood cell (WBC) counts from wild-type/control, Mfsd2b^-/-^, Spns2^-/-^, and ctDKO-Mx1Cre mice. WBC count from ctDKO-Mx1Cre mice was low as that of global knockout of Spns2. **F**, Deletion strategy of Mfsd2b and Spns2 in ctDKO-Mx1Cre mice induced by polyIC and the time points for S1P analysis. **G**, Plasma S1P level from control, ctDKO-Mx1Cre mice 1 month after polyIC injection, and Mfsd2b^-/-^ mice; (n=5, two experiments). Plasma S1P level from Mfsd2b^-/-^ mice was also included for comparison. **H-I**, Concentration (G) and percentage (H) of plasma S1P from 4 months old control, Mfsd2b^-/-^, Spns2^-/-^, and ctDKO-Mx1Cre mice. Plasma S1P from ctDKO-Mx1Cre mice was reduced by approximately 80%. Results shown in A, B, D, E, G, H and I are mean and SD, ****P<0.0001, **P<0.01, *P<0.05. One-way ANOVA was used.

### Additional source of plasma S1P from Mx1Cre-sensitive cells may exist independent of Mfsd2b and Spns2

To address whether there is an additional source of S1P that is independent of Mfsd2b and Spns2, we compared plasma S1P levels from ctDKO-Mx1Cre and SphK1^f/-^;SphK2^f/-^ Mx1Cre (hereafter: ckDKO-Mx1Cre) mice (**Figure 6A, B**). Our lipidomic results indeed showed that mice lacking both Mfsd2b and Spns2 still had higher levels of plasma S1P (~65% reduction) compared to that of SphK1 and SphK2 knockout mice (~98% reduction). These results suggest that an additional source of plasma S1P exists that is independent of Mfsd2b and Spns2 among Mx1Cre-sensitive cells (**Figure 6C**). To ascertain if plasma S1P can still be synthesized and released in ctDKO-Mx1Cre mice, we injected radioactive [3-^3^H] sphingosine (d18:1 sphingosine) into the circulation of ctDKO-Mx1Cre and control mice (**Figure 6D, E**). Then, the amount of [3-^3^H] S1P released into plasma and accumulated in RBC was analyzed within 2 hours post-injection. In control mice, [3-^3^H] S1P was detected in plasma within 10 minutes after [3-^3^H] sphingosine injection (**Figure 6D, E**) and gradually decreased over time (**Figure 6D**). In ctDKO-Mx1Cre mice, although significantly less plasma [3-^3^H] S1P was found 10 minutes after [3-^3^H] sphingosine injection, a detectable level of S1P was still observed in the plasma of ctDKO-Mx1Cre mice (**Figure 6D**). This novel source of S1P is independent from RBC as [3-^3^H] S1P was accumulated in RBC from ctDKO-Mx1Cre mice, whereas it was secreted from WT RBC over time (**Figure 6E**). Plasma S1P production is increased in response to irradiation or chemotherapy. We tested whether irradiation would increase plasma S1P in ctDKO-Mx1Cre mice. We found that sub-lethal dose of irradiation (6.5Gy) significantly increased plasma S1P production in control mice (SphK1^f/-^;SphK2^f/-^ or Mfsd2b^f/f^;Spns2^f/f^, without Mx1Cre) (**Figure 6F, G**). Although irradiation increased plasma S1P was largely dependent on Mfsd2b and Spns2, quantitatively a small amount of plasma S1P was increased in ctDKO-Mx1Cre mice (0.29μM v.s. 0.41μM), but not in ckDKO-Mx1Cre mice (0.022μM v.s. 0.046μM) (**Figure 6F, G**). These results indicate that ctDKO-Mx1Cre mice retain the capacity to release a minor amount of S1P.

**Figure 6.**
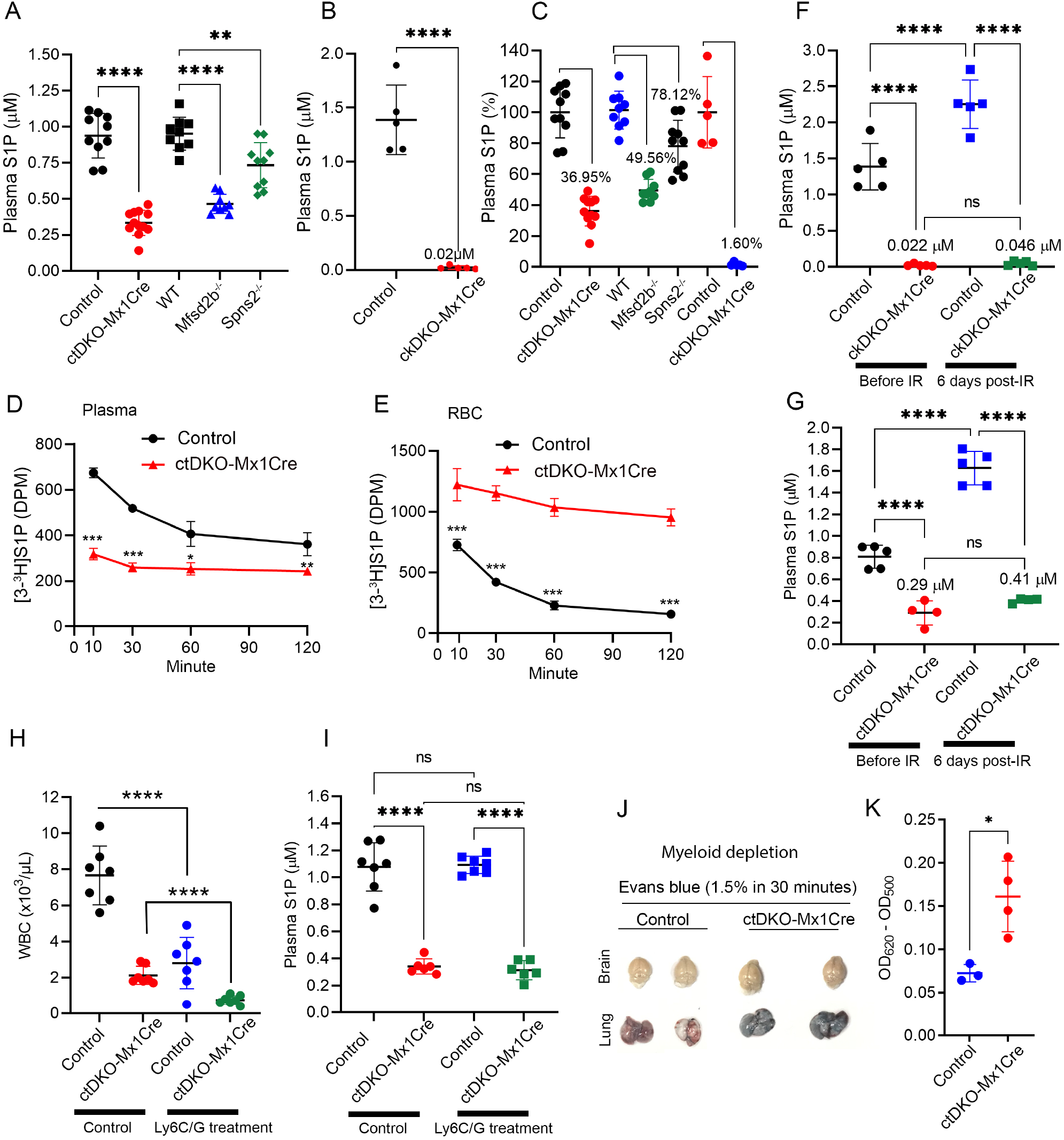
A minor source of plasma S1P independent of Mfsd2b and Spns2 may exist. **A-B**, comparison of plasma S1P levels from ctDKO-Mx1Cre and ckDKO-Mx1Cre (SphK1^f/-^, SphK2^f/-^ Mx1Cre) mice with their respective controls. Data are mean and SD. Each symbol represents one mouse; n=4-5 per genotype. **C**, Percentage of plasma S1P from indicated genotypes. Each symbol represents one mouse; n=4-5 per genotype. Results are mean ± SD, ****P<0.0001, ***P<0.001, *P<0.05. One-way ANOVA was used in A. t-test was used in B. **D-E**, In vivo S1P transport assay from control and ctDKO-Mx1Cre mice. [3-^3^H] S1P levels in plasma (D) and erythrocytes (E) of ctDKO-Mx1Cre and control mice at 10, 30, 60, and 120 minutes post-injection of [3-^3^H] sphingosine; n=4 per genotype from two experiments. Results are mean ± SD; ***P<0.001, **P<0.01, *P<0.05. Two-way ANOVA was used in D and E. **F-G**, Plasma S1P levels from controls, ckDKO-Mx1Cre, and ctDKO-Mx1Cre mice before (day 0) and 6 days post-irradiation with 6.5Gy. Note that the samples before irradiation in F and G was the same as in A and B. Each symbol represents one mouse; n=4-5 per genotype. Results are mean ± SD; ****P<0.0001, ***P<0.001, ns, not significant. One-way ANOVA was used in F and G. **H-I**, Depletion of myeloid cells from control and ctDKO-Mx1Cre mice. WBC count was reduced in both control and ctDKO-Mx1Cre mice after 40h treatment with Ly6C/G antibody. Plasma S1P levels in control or ctDKOMx1Cre mice treated with Ly6C/G antibody were comparable to the same genotype without treatment. Each symbol represents one mouse; n=6-7 per genotype. Results are mean ± SD, ****P<0.0001, *P<0.05, ns, not significant. t-test and One-way ANOVA was used in H and I, respectively. **J-K**, Accumulation of Evans Blue in the lungs of control and ctDKO-Mx1Cre mice 40 hours after treatment with Ly6C/G antibody. The lungs, but not the brains ctDKO-Mx1Cre mice were slightly more permeable with Evans Blue, but was not elevated compared to normal condition (See Figure 7B). Results are mean ± SD, *P<0.05.

Recently, monocytes were reported to provide S1P in lymph nodes in response to inflammation (Baeyens et al., 2021). We explored the possibility that the novel source of S1P is released from myeloid cells. Thus, we depleted myeloid cells including neutrophils and monocytes using Ly6C/G antibody (**Figure 6H**). Mice treated with Ly6C/G antibody had significantly lower WBC counts, indicating that the treatment was effective (**Figure 6H**). However, the levels of plasma S1P from ctDKO-Mx1Cre and control mice before and after Ly6C/G treatment were not reduced (**Figure 6I**). ctDKO-Mx1Cre mice had slightly basal leakage of Evans blue in the lung (see below, **Figure 8A, B).** Evans Blue leakage in the lungs of these myeloid-depleted ctDKO-Mx1Cre mice was not further elevated compared to the normal conditions (**Figure 6J, K**). Collectively, our results reveal that although Mfsd2b and Spns2 expressing cells provide most of plasma S1P, a minor amount of the lipid mediator is provided by other Mx1Ce-positive cells.

### Mfsd2b and Spns2 from Mx1Cre-sensitive cells provide essential sources of S1P for protection from anaphylactic shock

The significant reduction of plasma S1P of ctDKO-Mx1Cre mice prompted us to test whether the sources of S1P from Mfsd2b and Spns2 are required for protection from anaphylactic shock. Upon histamine challenge, core body temperature of control mice was decreased, but they had re-gained body temperature within 25-30 minutes, whereas core body temperature of ctDKO-Mx1Cre mice was decreased and remained low over 1 hour (**Figure 7A**). Approximately, 35% ctDKO-Mx1Cre mice died, whereas none of the control mice died during histamine challenge (**Figure 7B**). Blood of ctDKO-Mx1Cre mice had slightly higher hematocrit and significantly higher amounts of histamine in plasma, reflecting plasma loss and impaired glomerular filtration (**Figure 7C, D**). Their systolic blood pressure under that histamine challenge was significantly decreased (**Figure 7E**). Additionally, ctDKO-Mx1Cre mice were more susceptible to platelet activating factor (PAF) treatment (**Figure 7F**). To confirm if the significant reduction of plasma S1P in ctDKO-Mx1Cre mice is linked to the susceptibility to anaphylaxis, we performed rescue experiments in which exogenous S1P was intravenously injected to ctDKO-Mx1Cre and control mice during histamine challenge. Indeed, a bolus injection of S1P rescued core body temperature decrease from both control and ctDKO-Mx1Cre mice, whereas vehicle treatment did not improve their low body temperature induced by histamine (**Figure 7G**). These data indicate that circulating S1P is required to maintain blood pressure and that Mfsd2b and Spns2 provide a critical pool of S1P to protect mice from hypotension induced by histamine challenge. Together, our results indicate that the proportion of plasma S1P provided by Mfsd2b and Spns2 is essential for maintenance of vascular integrity and functions.

**Figure 7.**
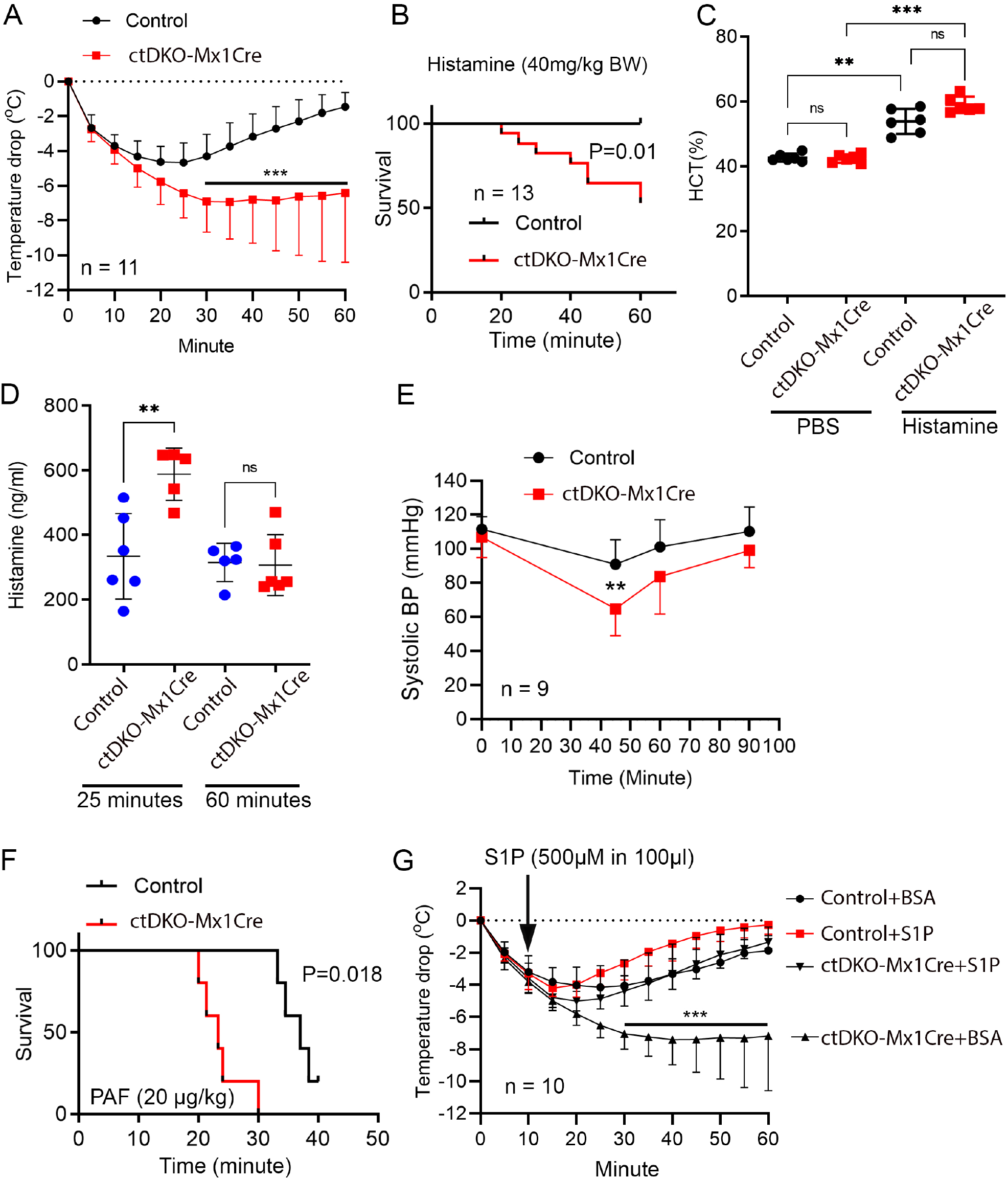
The sources of circulating S1P from Mfsd2b and Spns2 are required for protection from anaphylactic shock. **A,** Core body temperature drop was induced by intravenous injection of histamine (1mg/25g BW) into control and ctDKO-Mx1Cre mice (n=11; 3-4 months old). Core body temperature was monitored with a rectal probe. Results are mean ± SD, ***p<0.001. **B,** Survival curve of male control and ctDKO-Mx1Cre mice during histamine challenge (1mg/25g BW; i.v. route). Seven out of 20 ctDKO-Mx1Cre mice died, whereas all control mice survived (n=13 for WT, n=20 for ctDKO-Mx1Cre mice from 4 experiments). **C**, Hematocrit (HCT) of control and ctDKO-Mx1Cre mice measured after 25 minutes with histamine challenge (n=6 per genotype from 2 experiments). Results are mean ± SD, ***p<0.001, **p<0.01. **D**, Histamine levels in plasma of control and ctDKO-Mx1Cre mice 25 and 60 minutes after intravenous injection of histamine. Each symbol represents one mouse (n=4-5 per genotype). Results are mean ± SD, **p<0.01; ns, not significant. **E,** Systolic blood pressure was monitored in control and ctDKO-Mx1Cre mice before (0 minute), 45, 60, and 90 minutes after histamine challenge (0.25mg/25g BW; i.v. route). Results are mean ± SD, **p<0.01. **F**, Survival curve of control and ctDKO-Mx1Cre mice after a bolus injection of platelet activating factor (PAF, 20μg/kg; i.v. route). ctDKO-Mx1Cre mice exhibited significant susceptibility to PAF. n=5 per genotype. **G**, Exogenous administration of S1P rescued core temperature drop during anaphylaxis in ctDKO-Mx1Cre mice (n=10 per genotype, four experiments). Results are mean ± SD, ***p<0.001.

### S1P released by Mfsd2b and Spns2 from Mx1Cre-sensitive cells is required for maintenance of peripheral, but not CNS vasculature

Sustained S1P signaling activation is required for protection of blood vessel integrity. Thus, we examined whether peripheral and CNS vasculature of ctDKO-Mx1Cre mice are permeable to Evans Blue, a small dye that binds to BSA once injected into the circulation. We showed that at basal condition lungs of ctDKO-Mx1Cre mice were slightly accumulated with a higher level of Evans Blue (**Figure 8A, B**). Lack of S1P sources from Mfsd2b and Spns2 caused hemorrhage in the developing brain (See **Figure 2**). In contrast, we found that ctDKO-Mx1Cre mice had rather intact CNS vasculature at the normal condition (**Figure 8A, B; Supplemental Figure 6A**). Vascular insults disrupt junctions between endothelial cells and increase transcellular permeability. To examine whether S1P from Mfsd2b and Spns2 is required for the blood-brain barrier integrity under histamine treatment, we intravenously injected Sulfo-NHS-Biotin, a small molecule tracer, to ctDKO-Mx1Cre and control mice under histamine challenge. However, we did not observe significant extravasation of Sulfo-NHS-Biotin into the brain parenchyma of ctDKO-Mx1Cre mice (**Figure 8C**). These results indicate that S1P released from Mfsd2b and Spns2 is dispensable for CNS vasculature at normal and anaphylactic shock. In contrast, histological examinations of the lungs in ctDKO-Mx1Cre mice after histamine challenge showed that pulmonary blood vessels of ctDKO-Mx1Cre mice were thickened, indicative of edema (**Figure 8D, arrowheads**). We also found that the airways of ctDKO-Mx1Cre mice were filled with red blood cells, indicating that lung vasculature of ctDKO-Mx1Cre mice was compromised under histamine treatment (**Figure 8D, arrows**). Similar vascular leakage phenotypes were also observed in the lung of ctDKO-Mx1Cre mice after PAF treatment (**Supplemental Figure 6B**). Thus, our results indicate that S1P released from Mfsd2b and Spns2 is required for the protection of peripheral, but dispensable for CNS vasculature.

**Figure 8.**
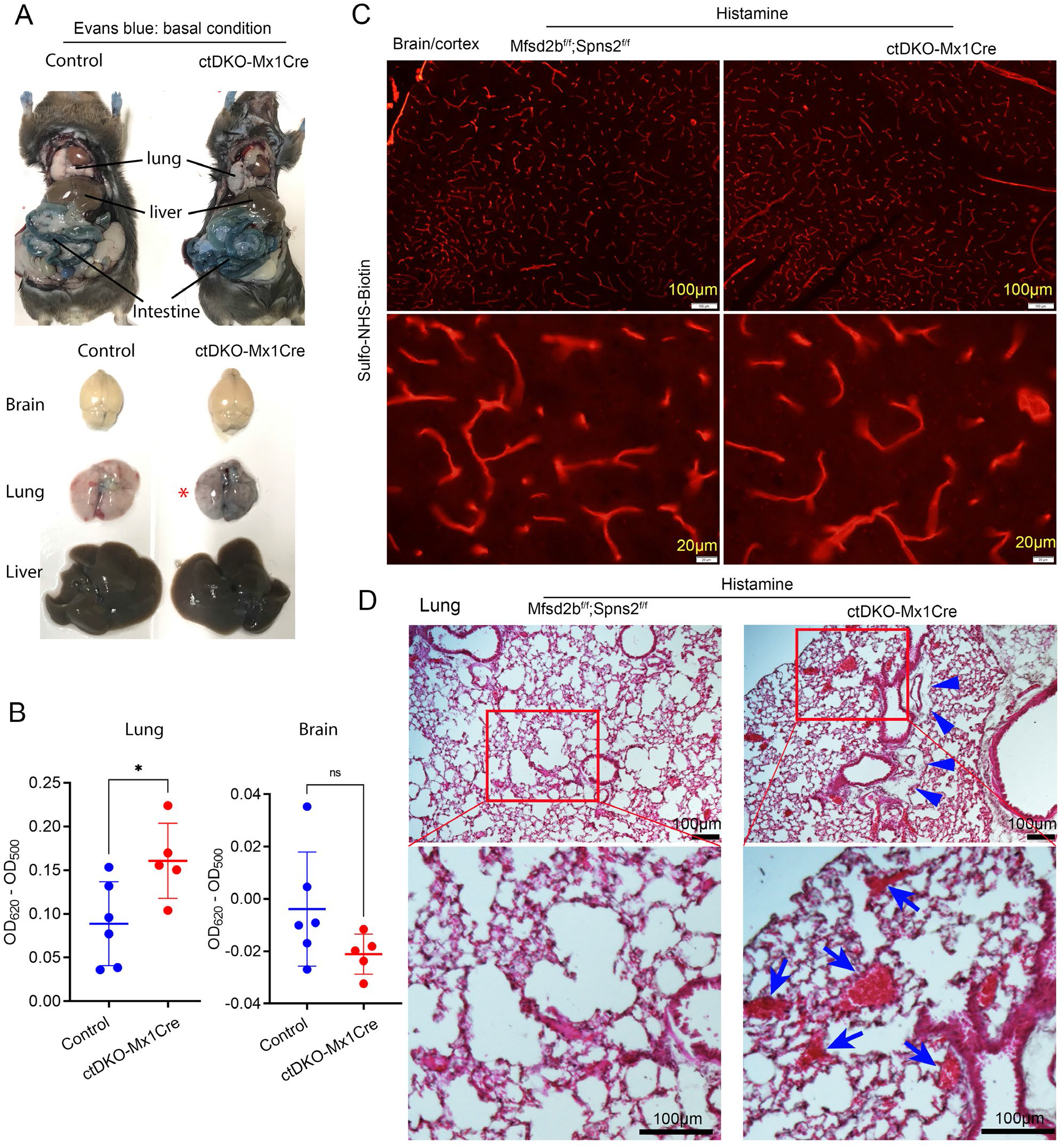
Compromised pulmonary, but not CNS vasculature of ctDKO-Mx1Cre mice. **A,** Evans Blue extravasation in the lung of control and ctDKO-Mx1Cre mice at normal condition showing that there was a slightly leakage in the lung of ctDKO-Mx1Cre mice (asterisk). **B,** Quantification of Evans Blue levels in the perfused lungs and brains from control and ctDKO-Mx1Cre mice. Each symbol represents on mouse. *p<0.05; ns, not significant; t-test. **C**, Examination of the blood-brain barrier leakage with Sulfo-NHS-Biotin in the cortex of control (Mfsd2b^f/f^, Spns2^f/f^) and ctDKO-Mx1Cre mice after histamine challenge. CNS blood vessels of ctDKO-Mx1Cre mice was not leaky to Sulfo-NHS-Biotin. Experiments were repeated twice (n=3). **D,** H&E staining of lung sections from control and ctDKO-Mx1Cre mice after histamine challenge (0.25mg/25g BW; i.v. route). Pulmonary blood vessels of ctDKO-Mx1Cre mice exhibited edema (arrowheads). Local infiltration of RBCs was also observed in the lungs of ctDKO-Mx1Cre mice after histamine challenge; n=6 per genotype from 3 experiments.

## Discussion

Sphingosine-1-phosphate (S1P) plays an essential signaling role in the cardiovascular system, particularly for maintaining blood vessel health. It is the extracellular ligand for 5 cognate G-protein coupled receptors (S1P1-5). The essential roles of S1P in the vasculature have been established via pharmacological or genetic perturbation of S1P receptors or sphingosine kinases (SphK1/2). Among the receptors, S1P1 is highly expressed in endothelial cells and plays a particularly important role in regulation of endothelial cell functions by sensing the circulating S1P. Mice lacking S1P1 or SphK1&2 are embryonically lethal due to severe hemorrhage (Liu et al., 2000), (Mizugishi et al., 2005). Postnatal ablation of S1P1 in the endothelial cells or compound conditional deletion of SphK1/2 also causes blood vessel leakage, especially in the pulmonary vasculature (Yanagida et al., 2017), (Camerer et al., 2009). The common view for S1P signaling is that S1P is synthesized inside cells and then exported to the extracellular milieu for signaling functions. Several cell types including endothelial cells and blood cells including erythrocytes can release S1P via the two recently identified S1P exporters Mfsd2b and Spns2. Interestingly, extracellular SphK1 was reported to generate S1P in the circulation. Adding to the complexity of S1P sources, several ABC transporters have been reported to exert the capacity for S1P transport (Mitra et al., 2006), (Kobayashi et al., 2009), (Vogt et al., 2018). To delineate the roles of Mfsd2b and Spns2, we generated global and conditional deletion of these genes. The results show that mice lacking both Mfsd2b and Spns2 die during gestation. Thus, these findings demonstrate that S1P from Mfsd2b and Spns2 is critical for embryonic development. Furthermore, we show in adult mice that, Mfsd2b and Spns2 are responsible for release of approximately 80% of the plasma S1P derived from SphK1&2 in Mx1Cre sensitive cells such as hematopoietic and endothelial cells. Intriguingly, our data also show that although most of the S1P is exported to plasma by Mfsd2b and Spns2, there may be an Mx1Cre-sensitive cell type that contributes 10-20% of plasma S1P that is independent of Mfsd2b and Spns2.

Sphingosine-1-phosphate (S1P) is synthesized in most cells in the body. However, only a few cell types are known to export S1P. Identifying the sources of S1P is critical to understand how it regulates immune and blood vessel functions. We have determined that Mfsd2b which is expressed in blood cells, contributes approximately half of plasma S1P (Vu et al., 2017). Among Mfsd2b-positive cells, erythrocytes and platelets were shown to provide plasma S1P (Chandrakanthan et al., 2021), (Nguyen et al., 2020), (Tan et al., 2020). In addition to S1P sources from Mfsd2b, S1P is also released via Spns2. The amount of plasma S1P contributed by Spns2 varies in different studies (Fukuhara et al., 2012), (Mendoza et al., 2012), (Nagahashi et al., 2013). In our analyses, we found that global deletion of Spns2 results in approximately 10-20% reduction of plasma S1P. This is recapitulated by the deletion of the transporter in endothelial cells (Mendoza et al., 2012). Nevertheless, lack of Spns2 causes severe lymphopenia suggesting that this transporter provides most of S1P in lymphatic vessels. These findings suggest that Spns2 provides specific sources of S1P, especially from lymphatic endothelial cells. However, it is unclear if Spns2 provides S1P during development. Our data show that embryos lacking Mfsd2b and Spns2 die between E13.4 and E14.5. The lethality to embryos lacking Mfsd2b and Spns2 is slightly delayed compared to the double knockout of sphingosine kinases, but largely recapitulates the phenotypes of S1P1 knockout. These results implicate that Mfsd2b and Spns2 coordinate to deliver S1P to S1P1 during development. Our results also argue against the roles of other transporters for S1P export during development, notably ATP-transporters that have been reported (Mitra et al., 2006), (Takabe et al., 2010), (Sato et al., 2007).

Global knockouts of S1P1 or SphK1&2 embryos exhibit severe hemorrhage and defects in the developing aorta. Consistently, embryos lacking Mfsd2b and Spns2 also have hemorrhage throughout the body, especially in lower dorsal regions. The main roles of S1P signaling in endothelial cells are to inhibit VEGF-induced tip cell formation and angiogenic hypersprouting, and stabilize adherens junctions. We observed that the compound transporter knockouts exhibited defective aortic structures in which endothelial cells were hyper-proliferating and the expression pattern of Claudin-5 was dysregulated. We did not observe significant reduction of the coverage of smooth muscle cells in the aorta. Instead, their coverage in the aortae was less compacted. These findings point to a downstream role of Mfsd2b and Spns2 in delivering circulating S1P from SphK1&2 to S1P1 for maintenance of the peripheral vasculature. In the developing brain, lack of Mfsd2b and Spns2 causes ruptured blood vessels in which the extravasation of red blood cells was seen in several brain regions. S1P receptors sense S1P at nanomolar concentrations and are expressed on both luminal and abluminal sides of endothelial cells. Deletion of Spns2, which is expressed in endothelial cells, is insufficient to cause these vascular phenotypes in the brain, implying that abluminal signaling of S1P is not critical for maintenance of CNS blood vessels. Mfsd2b is expressed in primitive red blood cells. Thus, it can only contribute to the circulating S1P. These findings imply that during development, Mfsd2b and Spns2 release S1P into the circulation, which is probably sensed by luminal S1P1 in endothelial cells for signaling functions. Recently, it was shown that Mfsd2a forms a complex with Spns2 to maintain the functions of the blood-brain barrier (Wang et al., 2020). Mfsd2a is expressed in the blood-brain barrier early during development (Nguyen et al., 2014), (Ben-Zvi et al., 2014). However, the presence of Mfsd2a is insufficient to rescue the lethality of Mfsd2b&Spns2 knockout embryos, arguing against its role as an S1P transporter.

Plasma S1P concentrations in adult human and mice are in a range of 1-2 micromolar, which exceeds the levels of S1P required for receptor activation (Kd=10-50nM) (Lee et al., 1998). Conditional deletion of S1P1 in endothelial cells results in vascular leakage in the lung as well as brain. Interestingly, mice deficient for plasma S1P by compound deletion of SphK1&2 only exhibit pulmonary leakage with Evans Blue (Gazit et al., 2016), (Camerer et al., 2009). In line with the results from SphK1&2 knockouts, the pulmonary vasculature of mice lacking Mfsd2b and Spns2 was also leaky to Evans Blue, but with lesser severity. In mice with compound loss of SphK1&2, the concentration of plasma S1P reduces by 98%, thus having approximately 20nM S1P in the circulation, while plasma S1P in the compound knockout of Mfsd2b and Spns2 mice is still >200nM. Interestingly, reduction of 50% plasma S1P in ApoM knockout mice is sufficient to cause pulmonary leakage of Evans Blue (Christensen et al., 2016). Thus, it seems that the concentration of S1P needed for activation of S1P receptors to protect peripheral vasculature is higher than shown in vitro (Lee et al., 1998). Perhaps, high levels of circulating S1P are also needed for activation of S1P receptors in perivascular cells for maintenance of vascular tone (Olivera et al., 2010). In contrast to the peripheral vasculature, cerebral blood vessels of adult mice lacking SphK1&2 or Mfsd2b&Spns2 are intact, suggesting that the remaining amount of S1P is sufficient to sustain the blood-brain barrier integrity or there is a novel source of S1P from neuronal cells that activates abluminal S1P1 receptor. This remains to be determined. Circulating S1P is required for protection of mice undergoing anaphylactic shock. We found that ctDKO-Mx1Cre mice had approximately 20% of normal plasma S1P and exhibited severe susceptibility to anaphylactic shock, indicating that high plasma S1P provided by Mfsd2b and Spns2 is essential to maintain vascular functions during pathological conditions. Supplementation of exogenous S1P to the circulation is sufficient to abolish the susceptibility to anaphylaxis in the knockout mice. Therefore, these findings emphasize the importance of having high plasma S1P concentrations that exceed the levels needed for S1P receptors’ activation.

Intriguingly, we found that compound knockout of Mfsd2b and Spns2 using Mx1Cre causes maximal reduction of 80% plasma S1P, whereas similar deletion of SphK1&2 reduces by 98% plasma S1P. These results suggest that there is a minor source of plasma S1P that is independent of Mfsd2b and Spns2 among Mx1Cre-sensitive cells. Deletion of SphK1&2 or Spns2 in endothelial cells only reduces 0-20% plasma S1P (Gazit et al., 2016), (Mendoza et al., 2012). This argues against the role of endothelial cells for the novel source of S1P. A recent study suggests that inflammatory monocytes secrete S1P during immune responses (Baeyens et al., 2021). Nevertheless, depletion of neutrophils and monocytes using Ly6C/G antibody failed to abolish the remaining amount of plasma S1P in the double Mfsd2b and Spns2 knockout mice, implying that other blood cells such as mast cells or eosinophils would be involved (**Supplemental Figure 7**). Identification of the cell types and the transporter responsible for this source of S1P should provide further understanding of how different sources of S1P regulate its receptor signaling.

Our study shows that Mfsd2b and Spns2 are the two major S1P transporters that provide S1P during embryonic development. These transporters from hematopoietic and endothelial cells provide a major pool of plasma S1P for maintenance of vascular functions, especially under pathological conditions. Interestingly, our results suggest that there may be another S1P transporter in Mx1Cre-sensitive cells, that provides approximately 20% of plasma S1P.

## Acknowledgements

This study was supported in part by Singapore Ministry of Health’s National Research Council NMRC/OFIRG/0066/20, Ministry of Education MOE2018-T2-1-126, NUSMED-FOS Joint Research Programme on Healthy Brain Aging grants (to L.N.N.) and the French National Research Agency (ANR-19-CE14-0028; to E.C.).

## Author Contributions

T.N.U.L., T.Q.N., Y.T.K.N., C.K.H.T., F.T., and P.K. performed in vivo experiments and ex vivo experiments. Z.H. and Y.T.K.N. performed H&E staining. L.C. performed and provided plasma S1P from SphK1/2 knockouts. E.C. designed experiments in SphK1/2 knockouts and edited the manuscript. M.R.W. and A.C-G. supervised lipidomic methodology. T.Q.N. performed lipidomic analysis. L.N.N. conceived and designed the study and experiments and wrote the paper. We thank Dr. Sangha Baik for initial work on the project.

## Competing interests

The authors declare no competing interests.

## Star Methods

### Mice

Global knockout mice of Mfsd2b were generated as described previously (Vu et al., 2017). Spns2 knockout mice were obtained from KOMP. All mice were in C57BL/6 background. Double global knockout (Mfsd2b^-/-^;Spns2^-/-^, hereafter gDKO) embryos were generated by inter-crossing Mfsd2b^-/-^;Spns2^+/-^ mice. Timed-mating was conducted; and pregnant Mfsd2b^-/-^;Spns2^+/-^ females were dissected at 12.5, 13.5, 14.5, and 15.5 days post-coitum to collect the fetuses. Double global knockout and control embryos (Mfsd2b^-/-^;Spns2^+/-^ or Mfsd2b^-/-^;Spns2^+/+^) were used. Double conditional knockout mice for Mfsd2b&Spns2 were generated by inter-crossing homozygous floxed Mfsd2b (Mfsd2b^f/f^) Spns2 (Spns2^f/f^) mice with Spns2^f/f^ Mfsd2b^f/f^ mice containing Mx1Cre. Deletion of both Mfsd2b and Spns2 was induced by polyIC. Three consecutive doses of polyIC (100μg/pub) were injected to intraperitoneal of P5 to P7 pups (P5-7). Compound conditional knockout Mfsd2b^f/f^;Spns2^f/f^;Mx1Cre (hereafter, ctDKO-Mx1Cre) with WBC of <2.5×10^3^/μl and control mice aged >3-12 months old (Mfsd2b^f/f^;Spns2^f/f^) were used for experiments. Deletion of Mfsd2b in red blood cells (RBC) was confirmed by S1P transport activity and Western blot, while deletion of Spns2 was confirmed by enumerating white blood cells (WBCs). Double conditional knockout mice for Sphk1&SphK2 (ckDKO-Mx1Cre) were generated by crossing Sphk1^f/f^:Sphk2^-/-^:Mx1Cre males to Sphk1^-/-^:Sphk2^f/f^ females and by induction of pups with polyIC postnatally as previously described (Gazit et al., 2016). Mice were maintained at a constant temperature of 20°C with 12-hour light/12-hour dark cycle on normal chow diets. All experimental protocols and procedures in the protocol R19-0567 were approved by IACUC committees under National University of Singapore.

### Chemicals

Sphingosine (D-erythro-sphingosine, 860490P-25MG), sphingosine-1-phosphate (D-erythro-sphingosine-1-phosphate, 860492P-1MG) were purchased from Avanti. Fatty acid-free bovine serum albumin (BSA) and histamine were purchased from Sigma (Cat. A7030-100G). PolyIC was purchased from Invitrogen (Cat. tlrl-pic-5). Sulfo-NHS-biotin was purchased from Thermo Fisher (Cat. 21217).

### Antibodies

The rabbit polyclonal antibody for Mfsd2b was generated as described previously (Vu et al., 2017) and used at 1:500 for Western blot. Gapdh was used as an endogenous control. Other primary antibodies and secondary antibodies used for immunofluorescent staining and Western blot are listed in table S2.

### Complete blood count

Blood was collected into EDTA-K2-coated tubes by heparinized capillary and directly counted using Celltex-α MEK-6400 (Nihon Kohden). Blood collected for WBC count was performed between 10am-12pm.

### Immunohistochemistry

Embryos were removed from the mother and fixed in 4% paraformaldehyde in PBS at 4°C overnight. They were then preserved in 30% sucrose (in water) for cryo-sectioning. Coronal cryo-sections of 30μm thickness from fetuses were prepared for immunohistochemistry with the following primary antibodies: anti-Glut1 (Abcam, Cat. ab40084), anti-Ter119 (BD pharmingen, Cat. 553670), anti-CD31(BD pharmingen, Cat. 553370), anti-CD144 (VE-Cadherin) (Invitrogen, Cat. 14-1441-82), anti-N-cadherin (Invitrogen, Cat. 33-3900), and anti-α-Smooth Muscle actin-Cy3™ (Sigma-Aldrich, Cat. C6918). Secondary antibodies used for detection included: AF488- (Invitrogen, Cat. A11006) and AF555- (Invitrogen, Cat. A21434). Sections were counterstained with 4,6-diamidino-2-phenylindole (DAPI) for cell nuclei visualization. Specimens were imaged with a confocal microscope (LSM710, Carl Zeiss) using 10x, 20x and 63x objectives.

### Transport assay with red blood cells

Blood was collected into 5% EDTA-K2 tubes and washed immediately twice with Tyrode-H buffer (10 mM HEPES-NaOH, 12 mM NaHCO_3_, 138 mM NaCl, 5.5 mM D-glucose, 2.9 mM KCl, 1 mM MgCl_2_, pH 7.4). For transport assays, 20 million erythrocytes (RBCs) were incubated with 2 μM [3-^3^H] sphingosine in 200 μl Tyrode-H buffer containing 0.5% BSA for 30 minutes at 37°C. Then, cell pellets (RBCs) and supernatants were separated by centrifugation at 1500 rpm for 5 minutes at room temperature. Cell pellets were washed once with 500 μl Tyrode-H buffer containing 0.5% BSA. Cell pellets and supernatant were subjected to S1P isolation using alkaline extraction. For radioactive counting, the same amount of aqueous phase containing S1P was mixed with scintillant (MP Biomedicals) and quantified using Tri-carb (GE Healthcare).

Similar transport assays were also performed with primitive RBC and fetal liver cells from Mfsd2b^-/-^ and Spns2^-/-^ embryos. For primitive RBC collection, E17.5 and E18.5 embryos were bled into warm PBS. Primitive RBCs were collected for transport assays as described above for mature RBC. Fetal livers from E17.5 and E18.5 embryos were also harvested and minced by a syringe plunger. The fetal liver cells were then passed through 75μm cell strainers before harvesting for transport assays.

### In vivo transport assay with [3-^3^H] sphingosine

Mice were intravenously injected with [3-^3^H] sphingosine solubilized in 12% BSA (Dose: 1μl of 1 mM [3-^3^H] sphingosine/gram body weight; specific activity: 10 μCi/ml). One minute later, 10μl blood was collected to monitor the initial signals. Blood samples were collected at 10, 30, 60, 120, and 180 minutes. Blood cells were spun down to collect plasma and RBCs. To harvest the remaining plasma, the cell pellets were washed once with Tyrode-H containing 0.1% BSA and spun down to collect the supernatant. The supernatant was combined with the plasma. The cell pellets and supernatant samples were subjected to S1P isolation by alkaline extraction. The radioactive signals of S1P from different time points were normalized to the S1P signal from WT collected at 10 minutes.

### Western blot analysis

Washed RBCs were lysed in RIPA buffer (25mM Tris pH 7.4, 150 mM NaCl, 0.1% SDS, 0.5% sodium deoxycholate, 0.5% Triton X-100) supplemented with protease inhibitor cocktail (Roche, 11836170001). An amount of 150 μg protein lysates was resolved on 10% SDS-PAGE, and transferred to nitrocellulose membrane. Brains were freshly harvested from E13.5 embryos and lyzed with RIPA buffer supplemented with protease inhibitor and phosphatase inhibitor cocktail 1 (Sigma-Aldrich, P2850). An amount of 75μg whole brain lysates was similarly utilized for Western blot analysis. Primary and secondary antibodies for detection are listed in table S2. ChemiDocMP (Bio-rads) system was used for detection. The protein bands were quantified by ImageLab software.

### Plasma collection for lipidomic analysis

Blood was collected under isoflurane anesthesia into EDTA-K2 collection tubes from the retro-orbital venous plexus with heparin-collected capillaries. Plasma was collected from blood samples after centrifugation at 2500 rpm for 30 minutes at room temperature. Blood samples with signs of hemolysis were excluded. Plasma was kept at −80°C until further analysis.

### Lipidomics

S1P analysis was performed using LC-MS/MS mass spectrometer as described previously (Narayanaswamy et al., 2014), (Chandrakanthan et al., 2021). Plasma samples from control and knockout mice aged 1-month and 4-months old were analyzed. For S1P analysis, plasma samples were first spiked with an internal standard (13C22H2-S1P, Toronto Chemicals), lipid extraction using a 1-butanol/methanol (1:1) mixture, followed by derivatisation with TMS-diazomethane. LC-MS/MS was performed on a UHPLC 1290 Agilent liquid chromatograph connected to a 6495A Triple Quadrupole mass spectrometer. Peak integration was performed using MassHunter Quantitative Analysis (QQQ) software, and data were manually curated to ensure correct peak integration. Areas under the curve (AUC) were corrected for isotopic interference where needed (Gao et al., 2021). One-point calibration was used to calculate the molar concentrations of the lipids. The stability of signal throughout the analysis was monitored by regular injection of a quality control samples. The levels of individual S1P were quantified and normalized to internal standards. Total levels of S1P were calculated and expressed as indicated in the figures.

### Body temperature measurements

Histamine (1mg/25g body weight) prepared in PBS was injected via retro-orbital vein to aged-match WT and ctDKO-Mx1Cre mice. Core body temperature of WT and ctDKO-Mx1Cre mice was recorded using a rectal probe with thermometer (RET-3, Type J/K/T Thermocouple Thermometer). Core body temperature was recorded every 5 minutes for a duration of 60 minutes after histamine injection. For rescue experiments, a bolus of S1P (18:1) in 12% BSA was i.v. injected to achieve final concentration of 500μM in blood. Injection volume was kept at 100μl for S1P and vehicle (12% BSA in PBS).

### Blood pressure measurements

Basal blood pressure was measured in both male WT and ctDKO-Mx1Cre mice aged 3-4 months using tail cup methods (BP2000, Visitech System, USA). Systolic blood pressure BP was monitored before histamine injection. For histamine-induced hypotension, histamine (0.25mg/25g BW) was injected via retro-orbital vein and systolic blood pressure was recorded 45, 60, and 90 minutes after histamine challenge. Six measurement cycles/each time point were collected and averaged.

### Histamine clearance

To determine the histamine clearance in plasma, histamine (1mg/25g BW) was intravenously injected to male ctDKO-Mx1Cre and control mice aged 3-4 months old. Blood was drawn at 25 and 60 minutes for plasma collection by centrifugation at 2500rpm as described above. Histamine levels were measured by ELISA according to vendor description (Bio Vison Histamine (HIS) Elisa kit; Cat. K4163-100).

### Sulfo-NHS-Biotin extravasation

WT and ctDKO-Mx1Cre mice were intravenously injected with histamine (1mg/25g BW). After 10 minutes, a bolus of Sulfo-NHS-Biotin (0.5mg/g body weight) was intravenously injected. Ten minutes later, mice were transcardially perfused with 10ml PBS for 5 minutes and the brain was fixed with 4% PFA for dissection. To detect biotin extravasation, streptavidin-Texas red was used as previously described (Wong et al., 2016), (Zhou and Nathans, 2014).

### Paraffin sections

For histological assessments, lungs were collected from 3-4 months old control and ctDKO-Mx1Cre mice 25 minutes after histamine challenge (1mg/25g BW, i.v. route). Lungs were immediately inflated with 1% PFA and fixed with 4% PFA for dissections. Paraffin sections from embryos were similarly prepared. For H&E staining, after dehydration using 70-100% ethanol followed by xylene (Sigma Aldrich, Germany, Cat. 1330-20-7), lung tissues or embryos were embedded in paraffin and then dissected using a microtome. The slides were tipped in xylene chamber to remove paraffin. Then, the slides were rehydrated progressively with 100 to 70% ethanol for 10 minutes each and rinsed with tap water for 5 minutes. The sections were stained with haematoxylin (Merck, Germany, Cat. 1.05174.2500) for 2 minutes and washed with tap water until excessed stain is thoroughly removed. Slides were tipped with 95% ethanol to dehydrate and then stained for 2 minutes with eosin (Sigma Aldrich, USA, Cat. HT110380). The excessed dyes were removed by washing with 50 to 100% ethanol in 2 minutes interval and the slides were cleared with xylene (Sigma Aldrich, Germany, Cat. 1330-20-7) for 5 minutes. The slides were air-dried and covered by DPX mountant (Merck, Germany, Cat. 06522) for imaging.

### Myeloid cell depletion

Control and ctDKO-Mx1Cre mice were injected via peritoneal with 500μg Ly6C/Ly6G antibody (BXcell, clone RB6-8C5, cat: BP0075). Ly6C/Ly6G antibody treatment depletes both monocytes and neutrophils. Complete blood cells were enumerated before and 40 hours after the antibody treatment. Plasma samples at these time points were also collected for S1P analysis as described above. Similarly, mice with myeloid depletion were also assessed for Evans Blue leakage. Control and ctDKO-Mx1Cre mice 40 hours post-treatment with Ly6C/Ly6G antibody were injected with 100μl of 1.5%Evans Blue in PBS (Sigma Aldrich, Germany, Cat. 314-13-6). The mice were perfused with 20 ml PBS for 5 minutes and lungs were harvested for quantification of Evans Blue accumulation.

### Evans blue extravasation

Control and ctDKO-Mx1Cre mice were injected with 100μl of 1.5% Evans Blue solution prepared in PBS through retro-orbital vein to assess vascular leakages in the lungs and brains. After 30 minutes, mice were anesthetized using ketamine solution (100 μl/10g BW, i.p. route) following by a perfusion with PBS solution. The lungs and brains were removed, photographed, and dried at 60° C overnight. To extract Evans Blue, dried lung and brain samples were immerged in 1 ml formamide solution (Sigma Aldrich, Germany, Cat. 75-12-7) for 4 days at room temperature. Evans Blue dye extract was determined as the absorbance of OD_620_ subtracted OD_500_.

### Irradiation

4-5 months old controls, ctDKO-Mx1Cre, and ckDKO-Mx1Cre mice were irradiated with a single dose of 6.5Gy. Plasma from the mice were collected before and 6 days post-irradiation for S1P analysis.

### Statistical analysis

Data was analyzed using GraphPrism9 software. Statistical significance was calculated using *t*-test or one and two-way ANOVA as indicated in the figure legends. P value < 0.05 was considered as statistically significant.

## Notes

### Competing Interest Statement

The authors have declared no competing interest.

