## Supplemental data for "Mfsd2b and Spns2 are essential for maintenance of blood vessels during development and protection of anaphylaxis"

### Supplemental figures 1-7

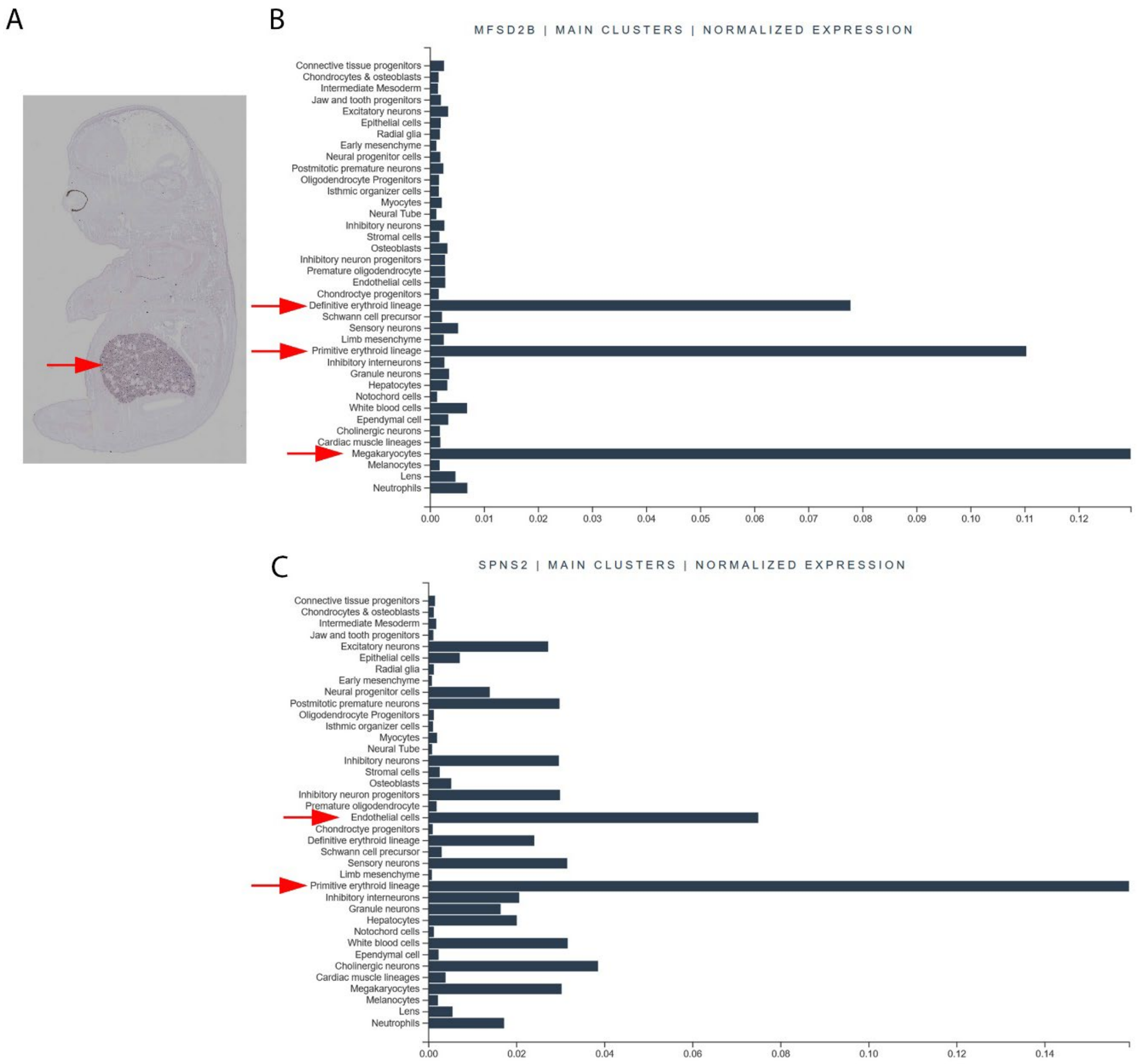


**Supplemental Figure 1. Expression of Mfsd2b and Spns2. A,** expression of Mfsd2b is found in fetal liver by in situ hybridization (arrow; source MGI:3583946). **B,** Expression of Mfsd2b is found in erythroid lineage including primitive and definitive red blood cells (arrows). **C,** Expression of Spns2 is found in endothelial cells and primitive red blood cells. Data in B and C were extracted from the Mouse Organogenesis Cell Atlas (MOCA): https://oncoscape.v3.sttrcancer.org/atlas.gs.washington.edu.mouse.rna/landing.

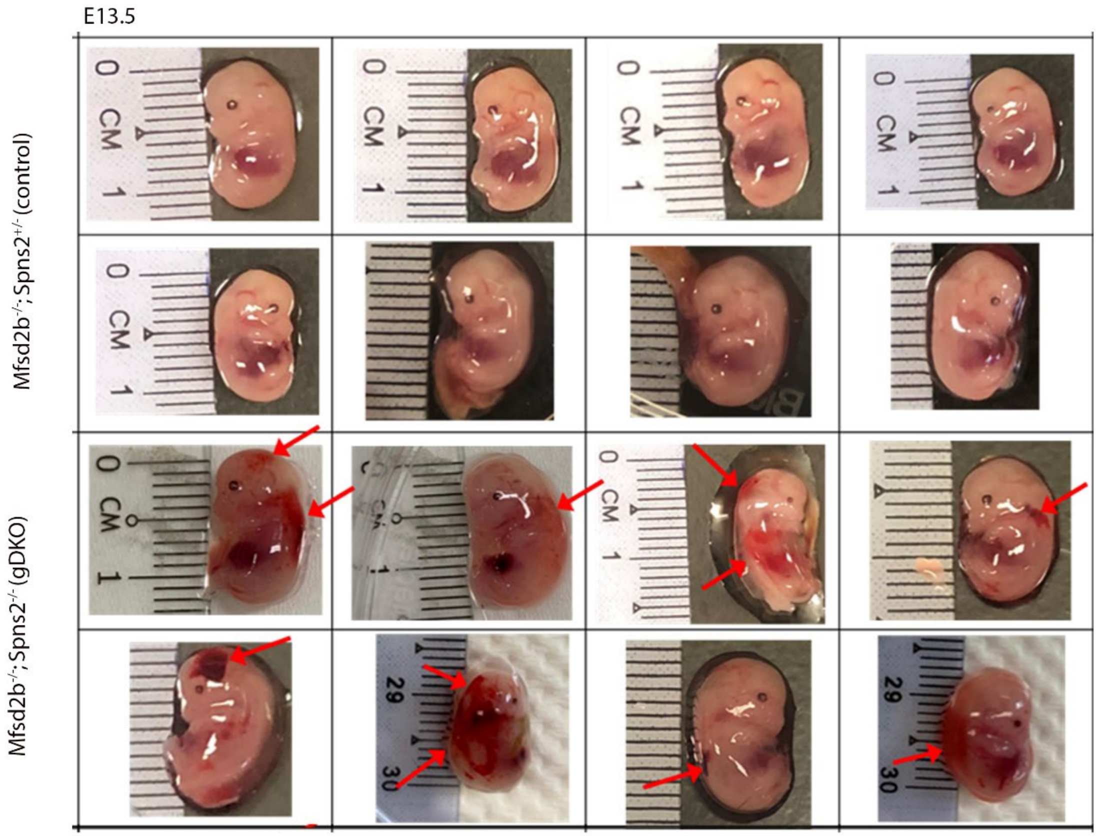


**Supplemental Figure 2. Deletion of Spns2 and Mfsd2b causes hemorrhage in dorsal and cranial regions**. In comparison with embryos carrying one Spns2 allele, the global knockout Mfsd2b and Spns2 embryos at E13.5 (gDKO) exhibited hemorrhage in the dorsal and cranial regions as indicated by the red arrows. n = 8 per genotype.

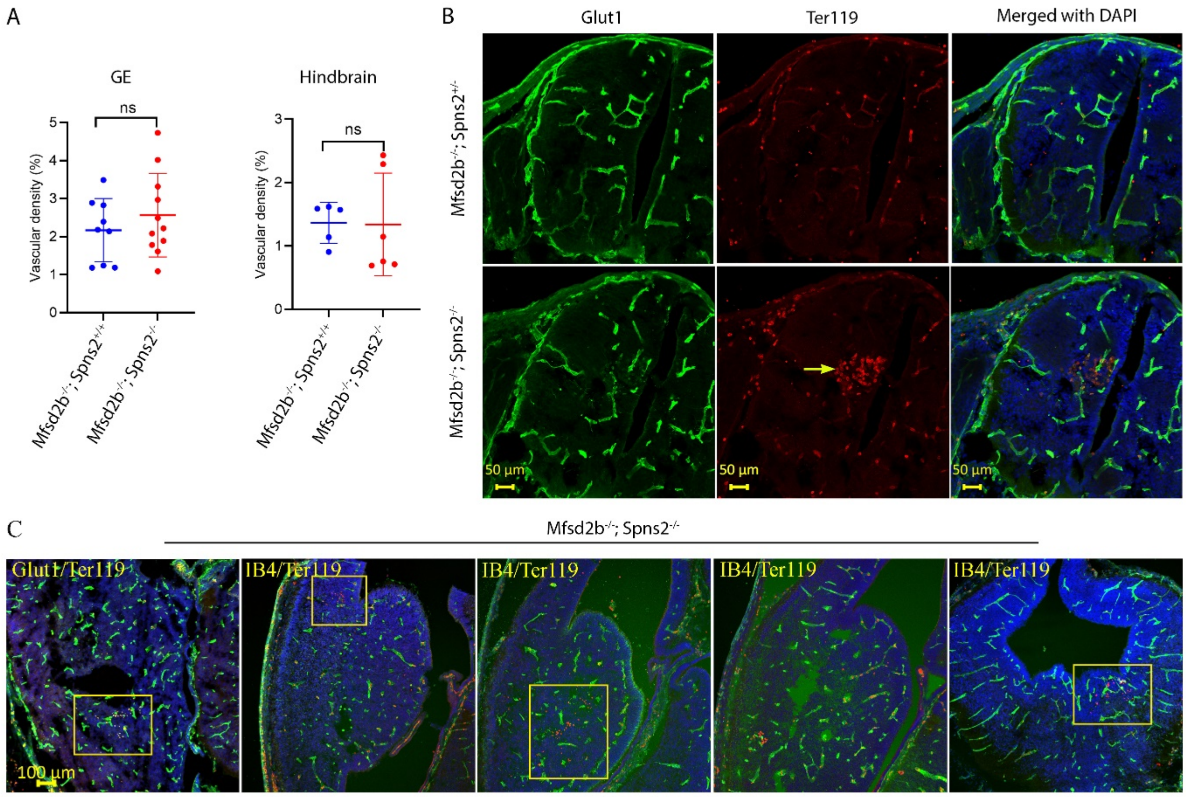


**Supplemental Figure 3. Compound global deletion of Spns2 and Mfsd2b causes local hemorrhage in the brain. A**, Vascular density in the ganglionic eminences (GE) and hindbrain. **B,** Immunofluorescence staining of control (Mfsd2b$^{-/-}$; Spns2$^{+/-}$) and gDKO embryos at E13.5 with endothelial cell marker Glut1 or Isolectin b4 (IB4) and erythrocyte marker Ter119. Hemorrhage was observed in the hindbrain/spinal cord of E13.5 gDKO embryos as indicated by the arrow. **C,** Hemorrhages in the GE regions of E13.5 gDKO embryos shown by yellow boxes. n = 5 for controls, n = 6 for gDKO.

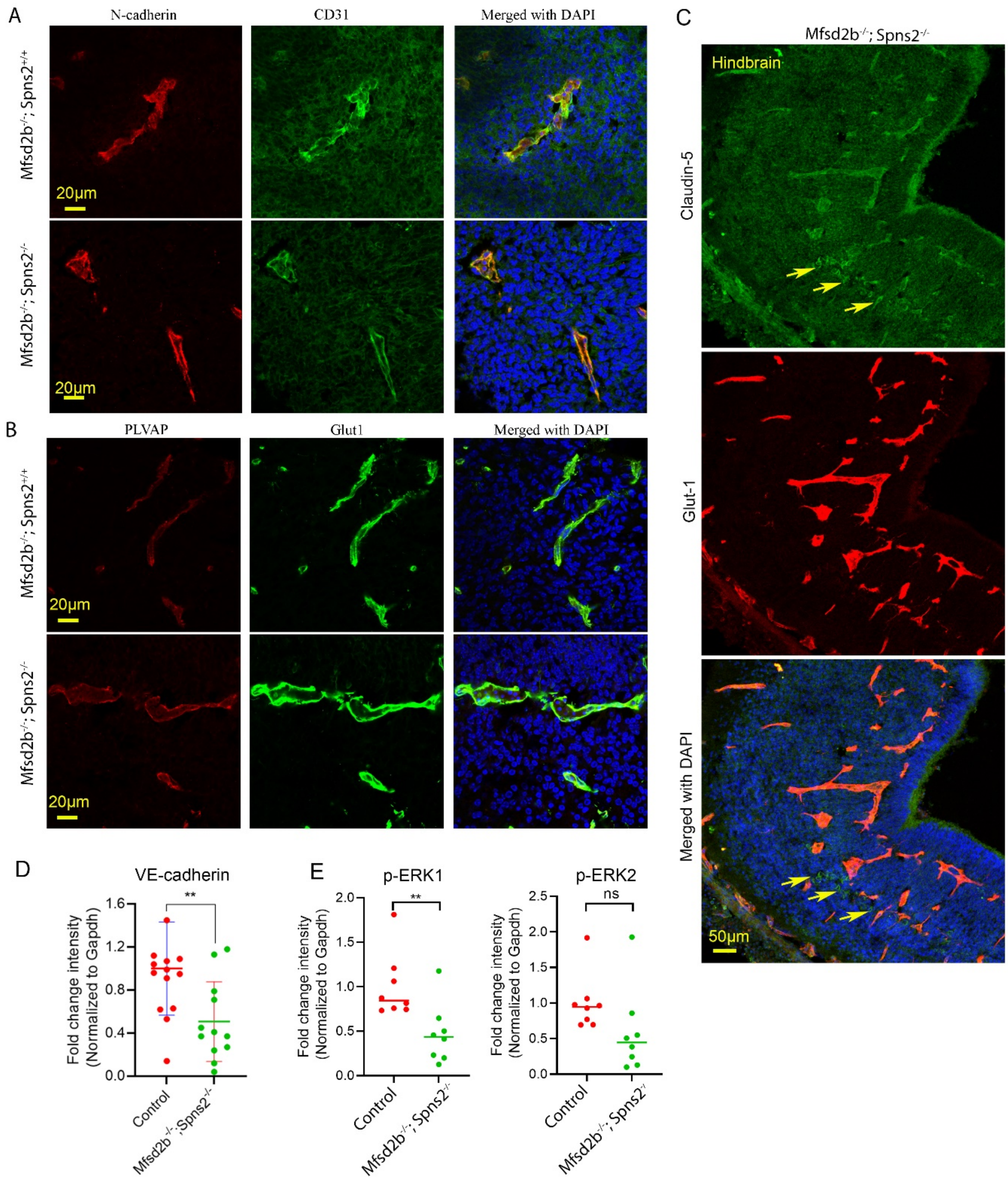


**Supplemental Figure 4. Lack of Mfsd2b and Spns2 affects endothelial tight junction proteins. A,** Localization of N-cadherin in CNS blood vessels of E13.5 control and gDKO embryos. CD31 was used to visualized blood vessels. Normal expression pattern of N-cadherin in CNS blood vessels of E13.5 gDKO embryos. **B,** Representative images of immunostaining of PLVAP, a fenestration marker, with Glut1 in the hindbrain of control and gDKO at E13.5. Glut1 was used to visualized blood vessels. PLVAP expression was not increased in CNS blood vessels of E13.5 gDKO embryos even in the hemorrhagic areas. n=3 per genotype. **C**, Localization pattern of Claudin-5 was dysregulated in the brain blood vessels of E13.5 gDKO embryos. Arrows show the ectopic expression of Claudin-5 outside of dilated blood vessels. **D**, Quantification of VE-cadherin expression band from Western blot analysis of the whole brain lysate of E13.5 controls (Mfsd2b$^{-/-}$; Spns2$^{+/+}$ and Mfsd2b$^{-/-}$; Spns2$^{+/-}$) and gDKO embryos. Each dot represents one mouse. Data are mean and SD. **P<0.01, t-test. **E**, quantification of phosphorylated ERK1/2 bands from Western blot analysis of the whole brain lysate of E13.5 controls (Mfsd2b$^{-/-}$; Spns2$^{+/+}$ and Mfsd2b$^{-/-}$; Spns2$^{+/-}$) and gDKO embryos. Experiments were repeated twice with n=2-3 per genotype. Data are mean and SD. **P<0.01, ns, not significant. t-test.

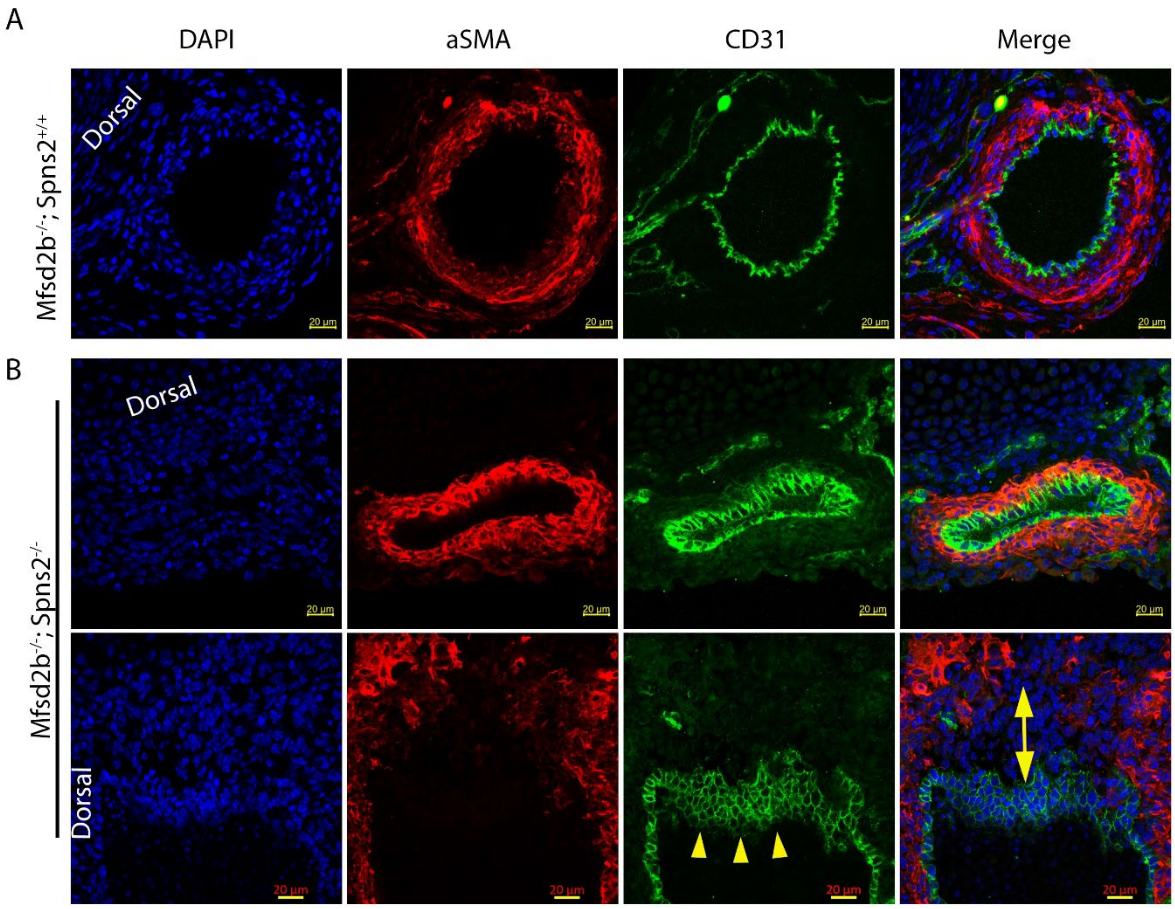


**Supplemental Figure 5. Defects in aortic structures of gDKO embryos. A-B,** deformation of aortic ring was observed in double knockout of Mfsd2b and Spns2 at E13.5. The endothelial cell layer shown by staining with CD31 (arrowheads) exhibited increased proliferation. The coverage of alpha-smooth muscle cells (aSMA) in the aortic ring of gDKO was also defective shown by the two-sided arrow. n = 9 for controls, n = 10 for gDKO.

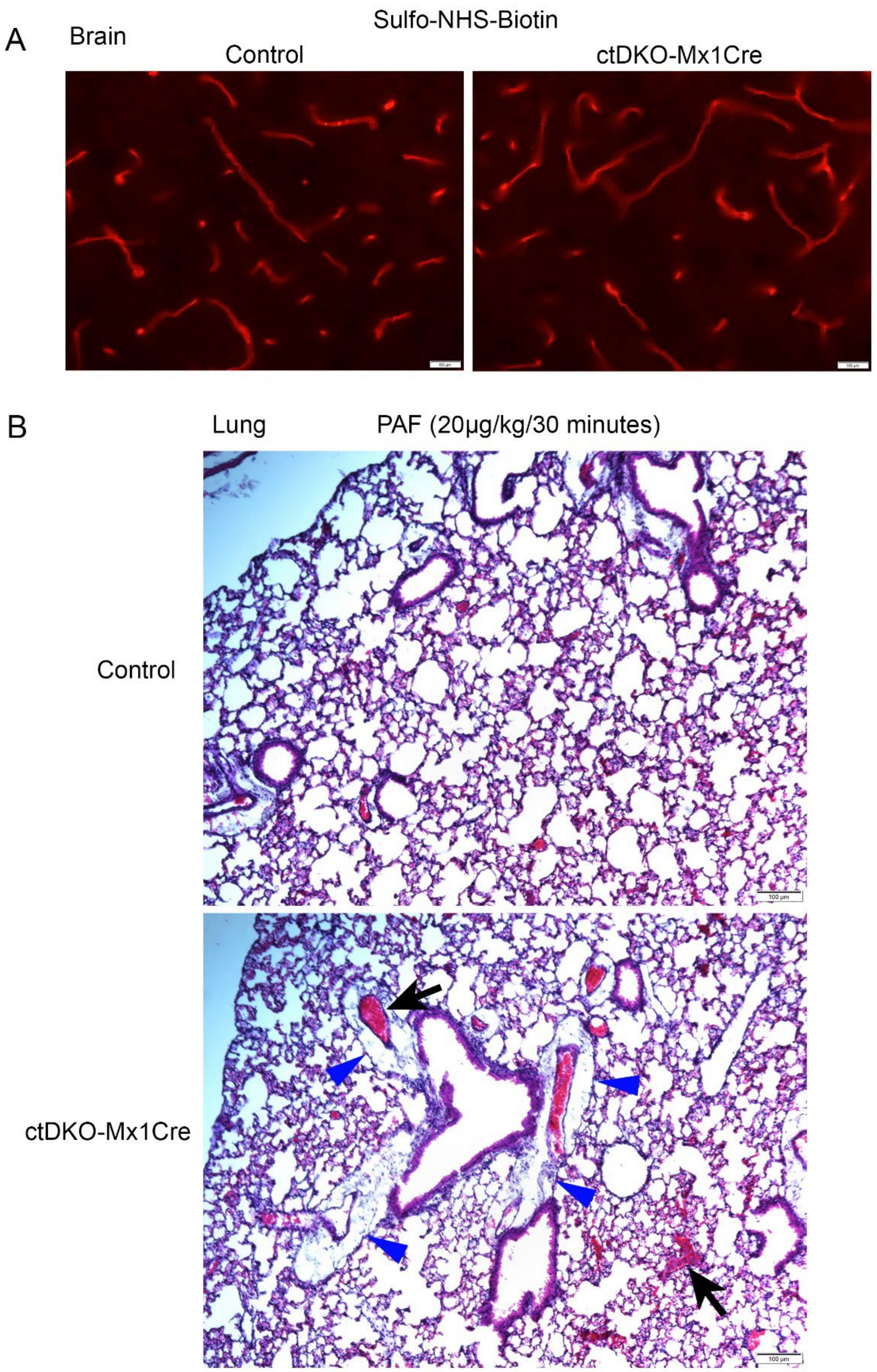


**Supplemental Figure 6. The lung, but not the brain vasculature of tDKO-Mx1Cre was compromised. A**, Normal blood-brain barrier functions of ctDKO-Mx1Cre mice. Shown are representative images of cortical brain sections after intravenous infusion with Sulfo-NHS-Biotin. There was no leakage of Sulfo-NHS-Biotin into the brain parenchyma of ctDKO-Mx1Cre mice. n=2 per genotype. **B,** Histology of lung from control and ctDKO-Mx1Cre after anaphylaxis induced by injection of PAF. Arrows show packed red cells in the lung blood vessels from ctDKO-Mx1Cre. Arrowheads show the signs of edema. n=2-3 per genotype.

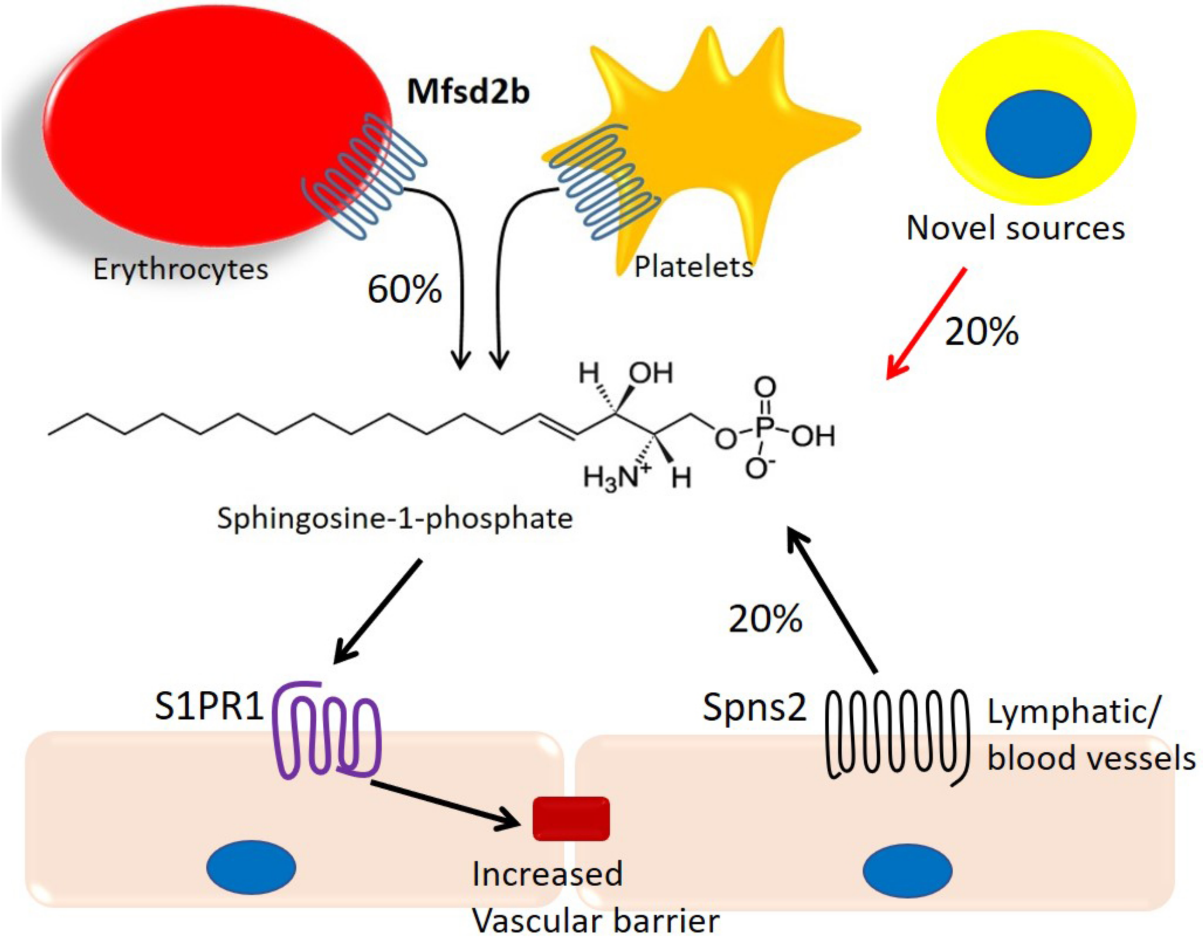


**Supplemental Figure 7. Schematic representation for various sources of plasma S1P**. The sources of S1P are critical to maintain vascular functions at the physiological and pathological conditions. Mx1Cre-sensitive cells including endothelial cells from lymphatic and blood vessels and hematopoietic cells generate most (>90%) of plasma S1P via the activity of sphingosine kinase 1 and 2. Among these Mx1Cre-sensitive cells, endothelial cells release approximately 20% S1P via Spns2, while erythrocytes and platelets supply 60% S1P via Mfsd2b. Global loss of Mfsd2b and Spns2 is lethal during development. However, compound deletion of Spns2 and Mfsd2b using Mx1Cre demonstrates that they together provide approximately 80% of plasma S1P. Although Mfsd2b and Spns2 provide essential sources of S1P, these results suggest that a novel transporter provides the remaining 20% plasma S1P.
